# The relationship between effective molarity and affinity governs rate enhancements in tethered kinase-substrate reactions

**DOI:** 10.1101/2020.03.12.989012

**Authors:** Elizabeth B. Speltz, Jesse G. Zalatan

## Abstract

Scaffold proteins are thought to accelerate protein phosphorylation reactions by tethering kinases and substrates together, but there is little quantitative data on their functional effects. To assess the contribution of tethering to kinase reactivity, we compared intramolecular and intermolecular kinase reactions in a minimal model system. We find that tethering can enhance reaction rates in a flexible tethered kinase system, and the magnitude of the effect is sensitive to the structure of the tether. The largest effective molarity we obtained was ∼0.08 µM, which is much lower than the effects observed in small molecule model systems and tethered protein-ligand interactions. We further demonstrate that the tethered, intramolecular reaction only makes a significant contribution to observed rates when the scaffolded complex assembles at concentrations below the effective molarity. These findings provide a quantitative framework that can be applied to understand endogenous protein scaffolds and to engineer synthetic networks.

## Introduction

Scaffold proteins assemble enzymes and their substrates into multiprotein complexes and are ubiquitous in signaling, metabolism, and protein homeostasis networks. While these proteins are widely assumed to promote reactions by increasing local concentrations, there is remarkably little quantitative data on the contribution of scaffold proteins to reaction rates. Nevertheless, cell biology and genetic studies clearly show that these proteins are critical for multiple biological functions.^1-3^ To understand how scaffold proteins affect reactivity in these biological processes, it is important to have well-defined biochemical models. Currently, however, we lack experimental systems to assess how large of a rate effect can be obtained from a scaffold protein, and how varying the structural properties of scaffold proteins affects reaction rates in protein assemblies. There is also substantial interest in engineering scaffold proteins for applications in synthetic biology and bioengineering.^4-8^ There have been notable successes in rewiring cell signaling networks with engineered scaffolds,^5-7^ but many attempts required screening large numbers of designs, and it is often unclear why plausible designs failed.^5,6,9-11^ Determining the criteria for effective protein scaffolds could enable more predictive rational design of scaffolded interactions.

To test the idea that proximity effects can increase reaction rates in kinase signaling reactions, and to develop a predictive framework for synthetic systems, we have constructed simplified model scaffolds using both covalent and non-covalent tethers that link a kinase to its substrate. We measured phosphorylation rates in both systems and compared them to the corresponding intermolecular, untethered kinase reaction (Figure 1). These comparisons allowed us to define an effective molarity for the intramolecular reaction (the ratio of the intramolecular to intermolecular rate constants), and to assess the effects of systematic variations of tether length on reactivity.

**Figure 1.**
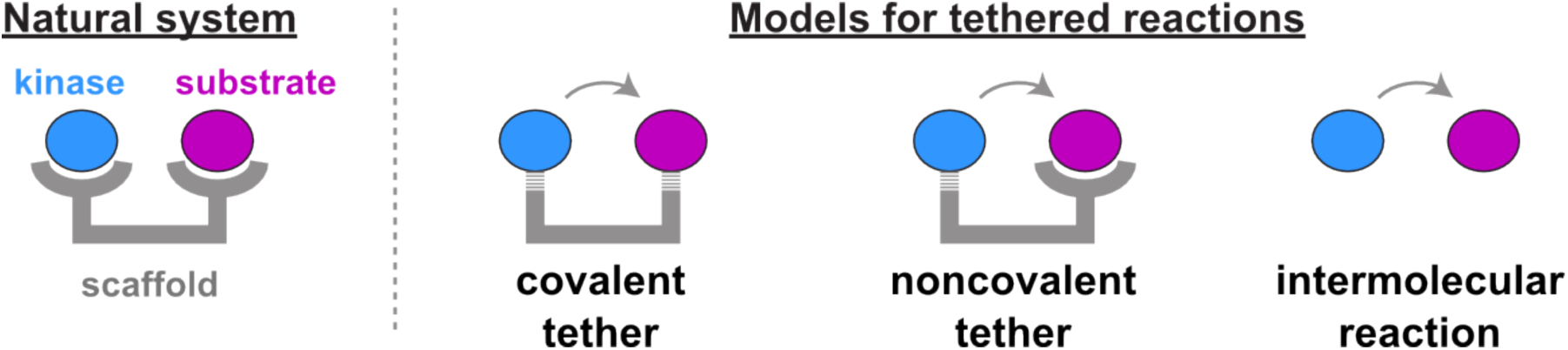
Models for scaffold-mediated kinase reactions. Natural scaffold proteins recruit kinases and their substrates (shown as blue and purple circles, respectively) to a scaffold (grey) using non-covalent protein interaction domains. To create a simplified scaffold model system, we recruited a kinase to a substrate using covalent and non-covalent tethers and compared them to an intermolecular reaction in the absence of scaffold.

Using this model system, we confirmed that a flexible tether can increase reaction rates, but the effects on kinase reactions are much smaller than expected compared to prior studies with small molecule and tethered protein-ligand model systems.^12-14^ Further experiments will be necessary to determine whether low effective molarities are general feature of enzymatic protein-protein reactions. We also demonstrate that the relationship between the affinity of complex assembly and the effective molarity is a critical parameter that governs rates in scaffolded reactions. When the binding affinity is weaker than the effective molarity, the scaffold complex does not assemble at substrate concentrations where the intramolecular reaction can proceed faster than the intermolecular reaction. These results provide a possible explanation for why simple peptide tethering strategies often fail in synthetic signaling pathways. More broadly, our work provides the foundation for a quantitative framework to evaluate the molecular function of protein scaffolds in natural assemblies and synthetic systems.

## Materials and Methods

### Protein expression constructs

All PKA (mouse residues 15-351, Uniprot P05132)^15^ and SYNZIP-fused substrate constructs were cloned into *E. coli* expression vectors containing an N-terminal maltose binding protein (MBP) and a C-terminal His_6_ tag. SpyCatcher-fused substrate constructs were cloned into *E. coli* expression vectors containing an N-terminal TEV-cleavable His_6_ tag.^16^ DNA for the SYNZIPs was kindly provided by Amy Keating (MIT) and DNA for SpyCatcher was kindly provided by Lauren Carter (University of Washington). We used a peptide derived from cystic fibrosis transmembrane conductance regulator (residues 693-705, FGEKRKNSILNPI) as a substrate.^17^ All linkers were composed of Gly, Ser, and Thr residues. These linkers are uncharged and predicted to be disordered.^14^ A complete list of protein expression constructs is provided in Table 1 and protein sequence information is provided in the Supporting Information.

**Table 1.** Protein expression plasmids.

| Plasmid | Vector Backbone<sup>a</sup> | Expressed Protein |
| --- | --- | --- |
| pES263 | pMBP-MG | MBP PKA-SpyTag H<sub>6</sub> |
| pES367 | pBH4 | H<sub>6</sub> Pep-2x-SpyCatcher |
| pES368 | pBH4 | H<sub>6</sub> Pep-4x-SpyCatcher |
| pES264 | pBH4 | H<sub>6</sub> Pep-8x-SpyCatcher |
| pES343 | pBH4 | H<sub>6</sub> Pep-14x-SpyCatcher |
| pES044 | pMBP-MG | MBP Pep-H<sub>6</sub> |
| pES234 | pMBP-MG | MBP PKA-SYNZIP6 H<sub>6</sub> |
| pES237 | pMBP-MG | MBP PKA-H<sub>6</sub> |
| pES311 | pMBP-MG | MBP Pep-SYNZIP5 H<sub>6</sub> |
| pES346 | pMBP-MG | MBP Pep-SYNZIP5<sub>trunc1</sub> H<sub>6</sub> |
| pES349 | pMBP-MG | MBP Pep-SYNZIP5<sub>trunc2</sub> H<sub>6</sub> |

<sup>a</sup> pMBP-MG is a modified version of pMAL-p2X (New England Biolabs) with an N-terminal TEV-cleavable MBP tag and a C-terminal His<sub>6</sub> tag. pBH4 is a modified version of pET15b (Novagen) with an N-terminal TEV-cleavable His<sub>6</sub> tag. pMBP-MG and pBH4 were described previously.<sup>34</sup> Complete protein sequences are available in the Supporting Information.

### Expression and purification of Synzip constructs

Proteins were expressed in Rosetta (DE3) pLysS *E. coli* by inducing with 0.5 mM IPTG overnight at 18 °C. All proteins were purified by Ni-NTA chromatography. Briefly, ∼5 mL cell culture pellet was resuspended in 25 mL lysis buffer (25 mM Tris, pH 8.0, 150 mM NaCl, 5 mM Imidazole, 2 mM MgCl_2_, 5 mM 2-mercaptoethanol, 5% glycerol, and containing one EDTA-free protease inhibitor tablet) and lysed by sonication. The subsequent lysate was cleared by centrifugation at 32,000xg for 50 minutes. Clarified lysate was incubated with Ni-NTA resin for 1 hour. The resin was washed four times with Buffer A (25 mM Tris, pH 8.0, 150 mM NaCl, 15 mM imidazole, 5% glycerol) and once with Buffer B (25 mM Tris, pH 8.0, 1 M NaCl, 5 mM imidazole, 5% glycerol). Protein was eluted using 25 mM Tris, pH 8.0, 150 mM NaCl, 250 mM imidazole, and 10% glycerol.

All proteins were dialyzed overnight into storage buffer (20 mM Tris, pH 8.0, 150 mM NaCl, 10% glycerol, and 2 mM DTT), aliquoted, and frozen at −80 °C. Protein concentrations were determined using a Bradford assay (Thermo Scientific).

### Expression and Purification of SpyTag-SpyCatcher constructs

MBP-SpyTag-H_6_ and H_6_-SpyCatcher proteins were expressed separately in Rosetta (DE3) pLysS *E. coli* and purified on Ni-NTA resin as described above. Prior to conjugation, proteins were dialyzed overnight into storage buffer at 4 °C to ensure complete removal of ATP. Subsequently, MBP-SpyTag-H_6_ and H_6_-SpyCatcher were mixed at a molar ratio of 1:10 in 1xPBS for 16-24 hours and the efficiency of the conjugation reaction was assessed by SDS-PAGE.

To remove excess H_6_-SpyCather from the reaction mixture, we further purified the SpyTag-Catcher fusion on amylose resin; the conjugate was applied directly to amylose resin and allowed to bind for at least two hours. The resin was washed five times with amylose wash buffer (20 mM Tris, pH 8.0, 200 mM NaCl, 2 mM 2-mercaptoethanol) and bound protein was eluted using amylose wash buffer that contained 15 mM maltose. Proteins were dialyzed overnight into storage buffer, aliquoted, and frozen at −80 °C.

### Kinetic assays

For SYNZIP-mediated non-covalent constructs, all kinetics measurements were performed in kinase assay buffer (40 mM Tris, pH 7.4, 200 mM NaCl, 10 mM MgCl_2_, and 0.05% IGEPAL). Enzyme concentration was held constant at either 1 nM or 2.5 nM and substrate concentrations varied over >20-fold range (from 0.01-0.25 µM). Phosphorylation was initiated by the addition of 100 µM ATP containing 0.02 µCi/µL γ-^32^P-ATP. This concentration of ATP is saturating in our system (Figure S11). Approximately 4.5 µL of the reaction was quenched at different time points by spotting onto nitrocellulose membrane and incubating in 0.5% phosphoric acid for at least 10 minutes. To remove excess ATP, the membranes were washed three to four times in 0.5% phosphoric acid, dried, and imaged on a GE Typhoon FLA 9000 imaging scanner. Kinetic parameters were determined using the initial rate method. In separate experiments, we varied the concentration of enzyme to verify that initial velocities scaled linearly with enzyme concentration (Figure S9).

Unimolecular rate constants (*k*_intra_) for SpyTag-mediated covalently tethered kinase-substrate complexes were measured using kinase assay buffer as described above. Intramolecular reaction rates were measured by varying the covalently-tethered complex concentration over >30-fold range (from 0.005-0.16 µM). At low [substrate], the resulting data (V_obs_ vs. [Enzyme]) was linear and the slope of this data corresponds to the intramolecular rate constant *k*_intra_ (Figure S2). The corresponding intermolecular reaction was measured using a fixed enzyme concentration (2.5 nM) and substrate concentrations that varied >200-fold range (from 0.02-5 µM). Initial rates were quantified as described above and the bimolecular rate constant (*k*_cat_/*K*_M_) was obtained from the slope of a linear fit to a plot of *k*_obs_ vs. [substrate]. In separate experiments, we varied the enzyme concentration to confirm that the observed rates for the intermolecular reaction were also first order in enzyme concentration (Figure S4).

To make the tethered PKA-product complex, we allowed the tethered PKA-substrate complex to react to completion. Briefly, we incubated 2.5 µM tethered PKA-substrate complex with 250 µM ATP for two hours at room temperature. To measure reaction rates, the fully phosphorylated complex was diluted to 2.5 nM and mixed with 100 µM cold ATP containing 0.02 µCi/ µL γ-^32^P-ATP. The reaction was initiated by the addition of free substrate. Initial rates were quantified as described above and the bimolecular rate constant (*k*_cat_/*K*_M_) was obtained from the slope of a linear fit to a plot of *k*_obs_ vs. [substrate].

### Competition Assay

For the competitive reaction with covalently-tethered PKA-substrate complex and free substrate, 0.02 µM covalently-tethered complex was mixed with free substrate that varied over a >200-fold concentration range (from 0.02-5 µM). At a given time point, 9 µL of the reaction was quenched with 6 µL 5xSDS dye. Subsequently, 10 µL of the quenched sample was analyzed by SDS-PAGE. After electrophoresis, gels were washed with 0.5% phosphoric acid twice for five minutes and imaged on a GE Typhoon FLA 9000 gel scanner.

### Fluorescence Polarization Binding Assays

Cys-containing proteins were labeled with Bodipy TMR using maleimide chemistry. 100 µM protein was reduced with 1 mM TCEP-HCl and incubated with at least a 10-fold molar excess of dye for 1 hour at room temperature. Free dye was removed by purification on a Ni-NTA column and/or a spin desalting column (Zeba). To determine labeling efficiency, proteins were separated by SDS-PAGE and subsequently imaged on a GE Typhoon FLA 9000 scanner at 532 nm.

1 nM labeled substrate was mixed with >700-fold range of unlabeled protein (from 1 to 700 nM) in binding buffer (20 mM Tris, pH 8.0, 150 mM NaCl, 2 mM TCEP, 2 mM MgCl_2_, and 0.05% IGEPAL). Reactions were incubated at room temperature for at least one hour in the dark. Polarization was measured in black 96-well plates using an EnVision 2105 multimode plate reader. The polarization of the peptide alone was subtracted from all measurements and the subsequent data was fit to a 1:1 binding model using equation 1:

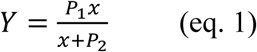

where *x* represents the protein concentration, *Y* represents the fluorescence polarization, *P*_1_ represents the maximum fluorescence polarization, and *P*_2_ is the dissociation constant.

Competition binding assays were conducted as described above using 1 nM labeled substrate, 30 nM PKA-SYNZIP6, and an unlabeled competing substrate that varied >10,000 fold range (from 1 nM to 13 μM). Background fluorescence polarization from non-specific binding between substrate competitor and labeled substrate was subtracted from each measurement. To determine the IC_50_, the plot of fluorescence polarization vs. [competitor] was fit to equation 2.

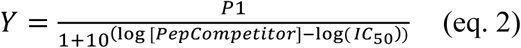

Subsequently, the IC_50_ was related to *K*_D_ using equation 3:

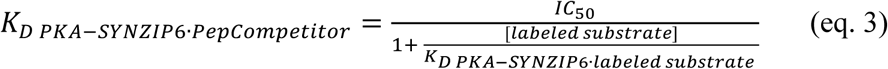

## Results and Discussion

We used the catalytic subunit of protein kinase A (PKA) as a model kinase because it has been extensively characterized both structurally and biophysically.^18-20^ Furthermore, PKA can be purified in active form from *E. coli* and can phosphorylate short peptides *in vitro.*^21^ We chose a natural PKA phosphosite derived from cystic fibrosis transmembrane conductance regulator (residues 693-705) as a substrate, which has a reported *K*_M_ of 30 µM.^17^ Thus, the reaction between PKA and the peptide substrate should not saturate with substrate concentrations in the nanomolar to low micromolar range (i.e. the reaction will be bimolecular), which provides a broad concentration range to assess tethering effects.

### Reaction rates for covalently tethered kinase-substrate complexes

To covalently tether PKA to a peptide substrate, we expressed PKA and its substrate separately and covalently fused them after purification via SpyTag and SpyCatcher proteins, which spontaneously form an irreversible isopeptide bond.^16^ This approach allowed us to avoid premature phosphorylation of the substrate, which would likely occur with a genetically-fused kinase-substrate complex. We constructed C-terminal fusions of PKA to SpyTag (PKA-SpyTag) via an eight residue Gly/Ser (GS) linker and the peptide substrate to SpyCatcher (Pep-SpyCatcher) via flexible linkers of variable length (Figure S1). After expression and purification, PKA-SpyTag and Pep-SpyCatcher constructs were mixed to form a covalently tethered kinase-substrate complex (Figure 2).

**Figure 2.**
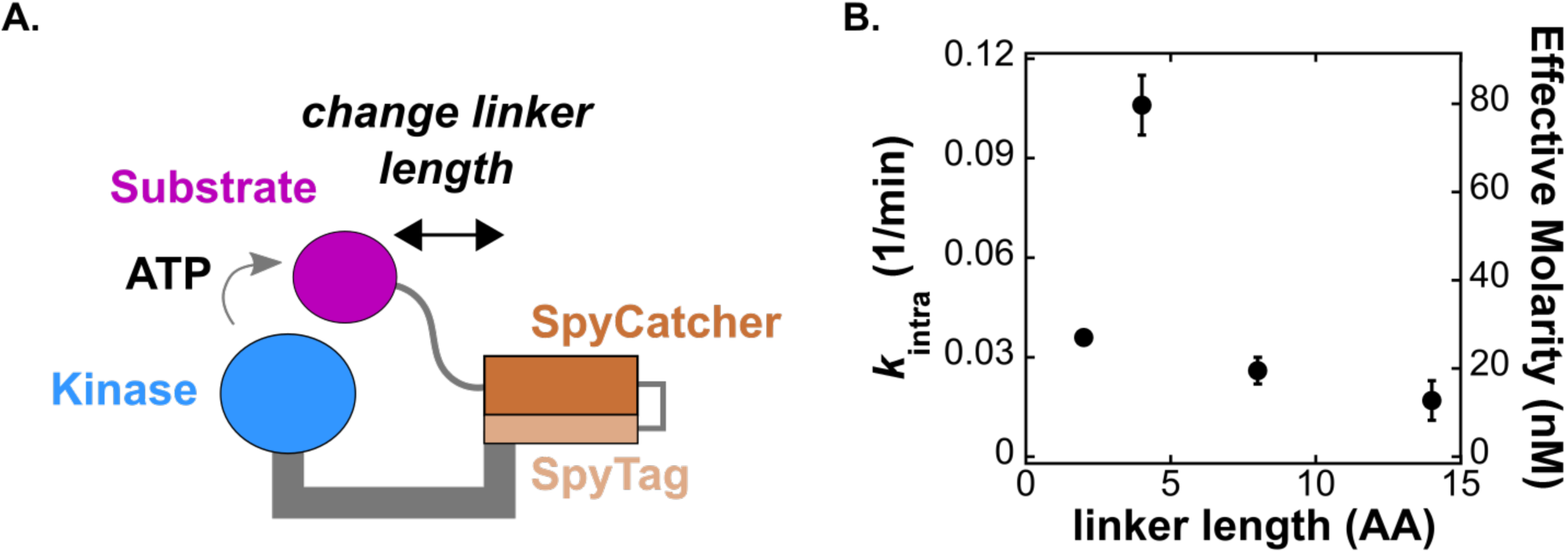
Reactivity depends on the structural properties of the assembly. (A) Schematic of the covalently tethered complex. The SpyCatcher-SpyTag complex covalently links the kinase with its substrate. We varied the length of the linker that connects the substrate to SpyCatcher from two to fourteen residues and measured the unimolecular rate constant (*k*_intra_). (B) Plot of *k*_intra_ vs. number of residues. The secondary y-axis shows the effective molarity for each complex, determined using the bimolecular rate constant from the untethered reaction (Figure 3A). Error bars represent the standard error of *k*_intra_ obtained from the fit of V_obs_ vs [tethered complex] as described in Figure S2.

With a covalently tethered kinase substrate complex in hand, we sought to compare reaction rates for tethered and untethered kinase-substrate reactions. Because tethered reaction rates are likely to depend on the length of a tether between the kinase and the substrate, we first measured substrate phosphorylation rates for a series of covalently tethered complexes with Pep-SpyCatcher linker lengths ranging from two to fourteen residues. For each complex, we measured initial rates of phosphorylation (V_obs_) using γ-^32^P-ATP to detect phosphorylated product formation and obtained the unimolecular rate constant (*k*_intra_) from a plot of V_obs_ vs. [tethered complex]. These plots are linear at concentrations below 50 nM, confirming that the observed reaction is intramolecular and not a bimolecular trans-phosphorylation reaction between distinct complexes (Figure S2). At concentrations above 50 nM, the observed rates increased non-linearly, likely due to competition from a bimolecular phosphorylation reaction. To extract *k*_intra_, we fit each dataset to a kinetic model that includes contributions from both the unimolecular and bimolecular reaction (Figure S2). We also obtained *k*_intra_ values by fitting the data below 50 nM to a linear model; these fitting methods produce *k*_intra_ values that are in close agreement (Figure S3).

As anticipated, we found that reaction rates depended on the structural properties of the tether. The observed values of *k*_intra_ vary over a 6-fold range with different Pep-SpyCatcher linker lengths and display a maximum of 0.11 min^-1^ with a four-residue linker (Figure 2 and Table 2). Thus, the four-residue linker appears to be sufficient to reach the kinase active site. The distance from the C-terminus of PKA to the C-terminus of a bound peptide is ∼23 Å (Figure S1),^15^ and this distance can plausibly be spanned by the four-residue linker because the construct also includes an eight-residue PKA-SpyTag linker. Pep-SpyCatcher linkers shorter and longer than four-residues are suboptimal. The shorter two-residue Pep-SpyCatcher linker may be unable to reach the active site of the kinase without distorting protein structure, while linkers longer than four residues likely result in a larger entropic barrier to finding the kinase active site.

**Table 2.**
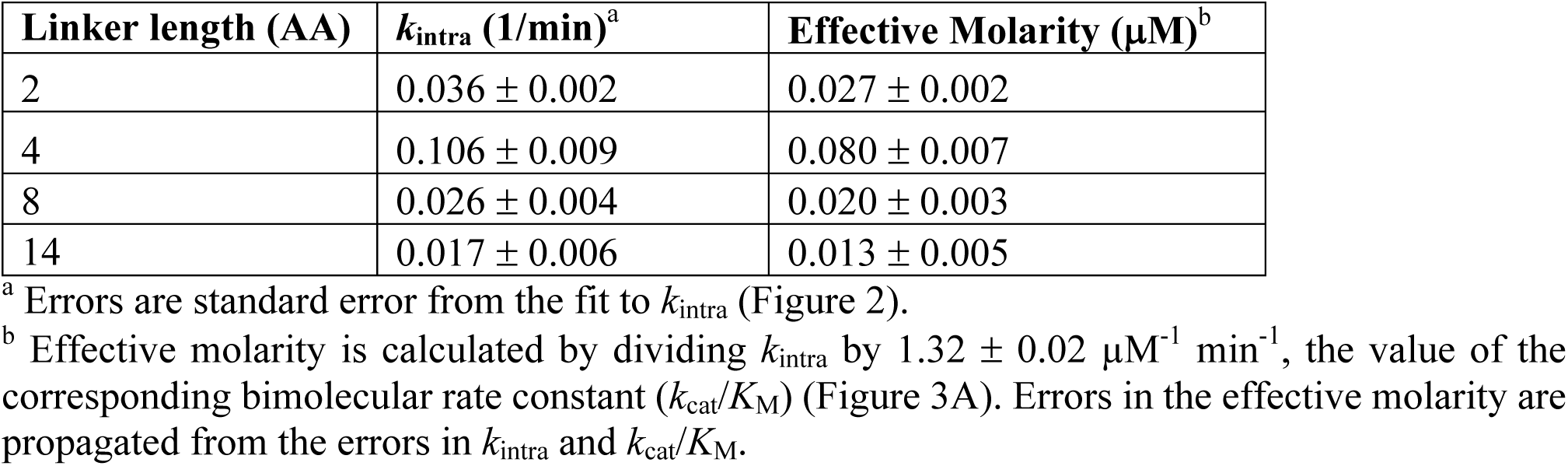
Kinetic parameters for covalently tethered complexes^a^.

| Linker length (AA) | $k_{\text{intra}}$ (1/min)<sup>a</sup> | Effective Molarity ($\mu\text{M}$)<sup>b</sup> |
| --- | --- | --- |
| 2 | $0.036 \pm 0.002$ | $0.027 \pm 0.002$ |
| 4 | $0.106 \pm 0.009$ | $0.080 \pm 0.007$ |
| 8 | $0.026 \pm 0.004$ | $0.020 \pm 0.003$ |
| 14 | $0.017 \pm 0.006$ | $0.013 \pm 0.005$ |

<sup>a</sup> Errors are standard error from the fit to $k_{\text{intra}}$ (Figure 2).
<sup>b</sup> Effective molarity is calculated by dividing $k_{\text{intra}}$ by $1.32 \pm 0.02 \mu\text{M}^{-1} \text{min}^{-1}$ , the value of the corresponding bimolecular rate constant ( $k_{\text{cat}}/K_{\text{M}}$ ) (Figure 3A). Errors in the effective molarity are propagated from the errors in $k_{\text{intra}}$ and $k_{\text{cat}}/K_{\text{M}}$ .

### Effective molarity for the covalently-tethered reaction

To identify the conditions in which covalent tethering can increase reaction rates, we compared the intramolecular reaction to an intermolecular reaction containing PKA-SpyTag with a free, untethered substrate. For the free reaction, we measured initial rates over a >200-fold range of substrate concentration (0.02 – 5.1 µM) and a 10-fold range of enzyme concentration (Figure 3A and Figure S4). In these concentration ranges, observed rates scaled linearly with substrate and enzyme concentration, indicating that the observed reaction is bimolecular. For this reaction, there was no detectable saturation at high substrate concentration, as expected from the reported *K*_M_ of 30 µM.^17^ We therefore fit *k*_obs_ (i.e. V_obs_/[E]) vs. [substrate] to a linear model, where *k*_obs_ = (*k*_cat_/*K*_M_)[S] and obtained a bimolecular rate constant (*k*_cat_/*K*_M_) of 1.3 µM^-1^ min^-1^.

**Figure 3.**
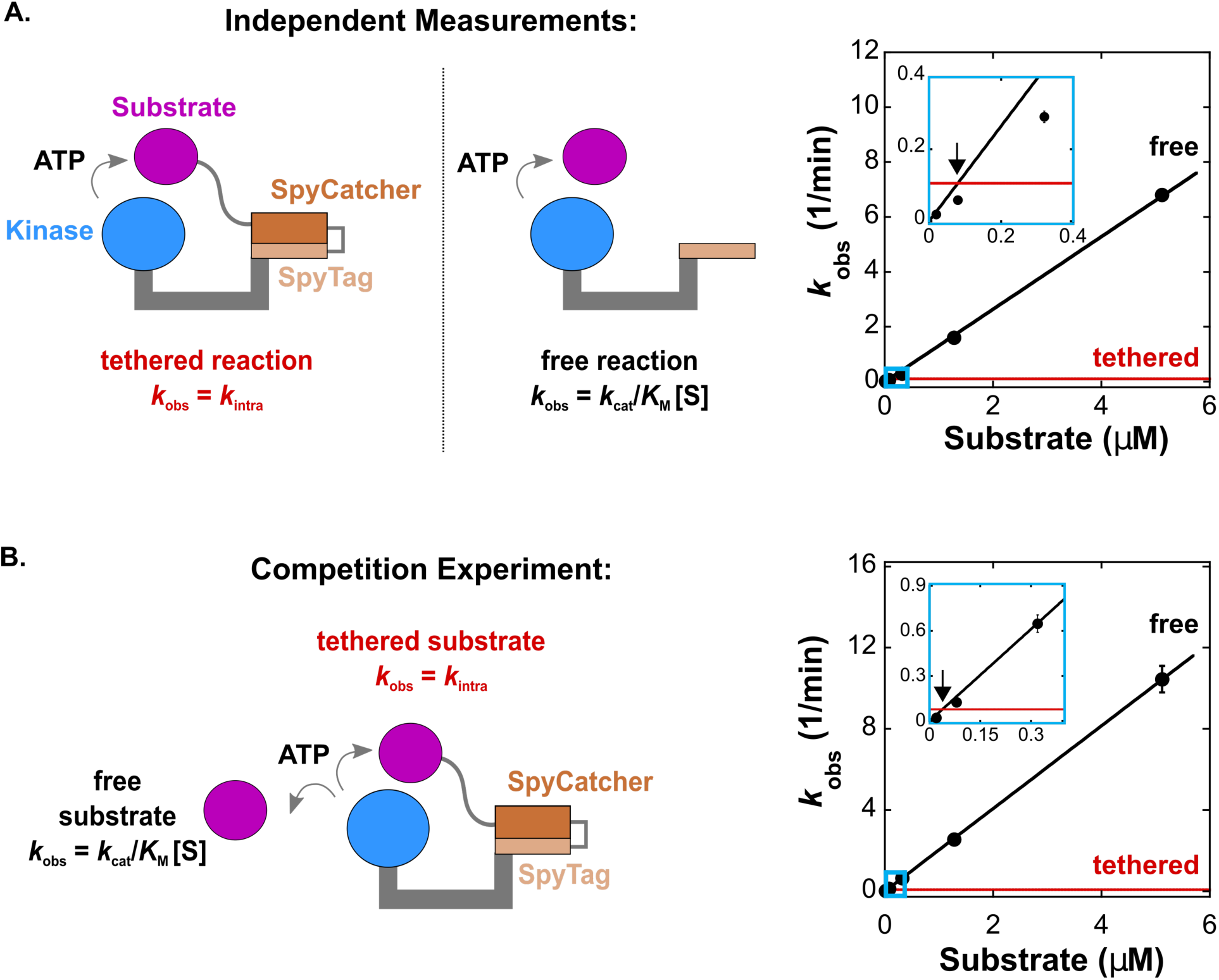
Intermolecular reactions readily outcompete the covalently tethered reaction. (A) Schematic of the tethered (4 aa linker) and free reaction and corresponding plot of *k*_obs_ vs. [substrate] for the tethered (red) and free (black) reactions. Inset is marked with light blue rectangle. The observed effective molarity is 0.08 µM. (B) Schematic of a competitive reaction with both tethered (4 aa linker) and free substrates present, and corresponding plot of *k*_obs_ vs. [substrate] for the tethered (red) and free (black) reactions. Inset is marked with light blue rectangle. The observed effective molarity is 0.04 µM. Each [product] vs time trace was measured from separate reactions in duplicate. For (A) and (B), error bars represent the standard error for *k*_obs_ obtained from a linear fit to [product] vs time for both datasets. The error for *k*_intra_ is smaller than the thickness of the red line (see Table 2).

The effective molarity is the substrate concentration at which the intermolecular and intramolecular rates are equal and can be obtained by finding the point at which the rate constant *k*_obs_ is equal for the two reactions. For the tethered reaction, *k*_obs_ = (*k*_cat_/*K*_M_)[S] and increases linearly with increasing [substrate]. For the free reaction, *k*_obs_ = *k*_intra_ and is constant. Plotting these values together gives a pair of lines that intersect at the effective molarity. For the optimal four-residue tethered complex, the observed effective molarity is 0.08 µM (Figure 3A). At substrate concentrations <0.08 µM, the intermolecular reaction is slower than the covalently-tethered reaction, while at substrate concentrations >0.08 µM the intermolecular reaction is faster. As with the observed value of *k*_intra_, the effective molarity displays a maximum with a four residue Pep-SpyCatcher linker (Figure 2B). Similar effective molarity trends with a maximum at an intermediate linker length have been observed in tethered protein-ligand model systems.^15^

To evaluate the significance of an effective molarity of 0.08 µM, we can make comparisons to other systems where these values have been measured. Small molecule reactions have effective molarities that can range above 10^8^ M,^12,22^ while tethered protein-ligand interactions have measured and predicted values in the mM range.^13,14,23^ Further, we can obtain a rough estimate of an effective concentration for a tethered substrate by using the covalent tether length to define a spherical volume where the substrate is confined (Figure S5), which gives a value of ∼4 mM. Based on these comparisons, the observed effective molarity of 0.08 µM for our tethered kinase-substrate system is unexpectedly small. However, effective molarity is a complex parameter that also depends on orientation effects from rotational and vibrational degrees of freedom.^22^ There is limited data available for effective molarities in tethered enzymatic protein-protein reactions, and it remains to be seen whether there is a general difference in behavior between tethered enzymatic reactions and binding interactions. To obtain larger effective molarities in kinase-substrate reactions, it may be necessary for a scaffold to precisely constrain the orientation of the substrate relative to the active site.

### The covalently-tethered system does not form a stable product-bound complex

One potential tradeoff associated with scaffold-mediated rate enhancements is that, by promoting interactions between a kinase and a substrate, the scaffold may stabilize the product-bound state and inhibit multiple turnovers of the enzyme. To test this possibility, we measured reaction rates in a competitive reaction between free substrate and substrate covalently tethered to enzyme. If product inhibition by the tethered product limits turnover, phosphorylation of the free peptide by the tethered PKA-substrate complex should be slower than the reaction of untethered substrate with free PKA.

To measure rates for a tethered and free substrate in the same reaction, we incubated the covalently tethered PKA-substrate complex with varying concentrations of free substrate and monitored phosphorylation of both substrates simultaneously using SDS-PAGE and autoradiography (Figure S6). We used 20 nM tethered complex, a concentration at which the bimolecular reaction between two tethered complexes is insignificant (Figure S2). From the observed rate of phosphorylation of the free substrate, we obtained a plot of *k*_obs_ vs. [substrate] and a bimolecular rate constant (*k*_cat_/*K*_M_) of 2 min^-1^ µM^-1^ (Figure 3B). In the same reaction, we measured the phosphorylation rate of the tethered substrate and used *k*_intra_ = V_obs_/[ES]_tethered_ to obtain a unimolecular rate constant of 0.08 min^-1^. The values of *k*_cat_/*K*_M_ and *k*_intra_ from this competition experiment are similar to our previous measurements of these values in separate reactions (1.3 min^-1^ µM^-1^ and 0.08 min^-1^, respectively) (compare Figure 3A and 3B). In particular, the identical *k*_intra_ value indicates the tethered complex is still functional for the intramolecular reaction in these conditions, and the absence of any decrease in *k*_cat_/*K*_M_ indicates that the tethered substrate does not stably occupy the active site and inhibit enzyme turnover.

To independently confirm that tethered product does not inhibit enzyme turnover, we also prepared a fully phosphorylated tethered complex by allowing the covalently tethered PKA-substrate complex to react to completion. When the tethered PKA-product complex is mixed with varying concentrations of free substrate, the observed bimolecular rate constant (*k*_cat_/*K*_M_) is 1.4 min^-1^ µM^-1^, indistinguishable from the *k*_cat_/*K*_M_ of 1.3 min^-1^ µM^-1^ for the free reaction (Figure S7).

Importantly, these results directly demonstrate the conditions in which an untethered reaction outcompetes the tethered reaction. The plot of *k*_obs_ vs. [substrate] gives a pair of lines that intersect at an effective molarity of 0.04 µM (Figure 3B), similar to the value of 0.08 µM obtained when the rates are measured in separate reactions (Figure 3A). At concentrations above the effective molarity, even when a substrate is covalently tethered to PKA, the free reaction outcompetes the tethered reaction.

### A non-covalent tether can accelerate the kinase reaction at low substrate concentrations

Natural scaffold proteins use non-covalent protein-protein interactions to recruit multiple signaling proteins to the macromolecular complex. To understand how protein-protein interactions affect tethering, we constructed a model system where the substrate can bind non-covalently to a remote binding site on the kinase (Figure 1). For simplicity, the kinase remains covalently fused to the scaffold in our system. This approach allows us to make a direct comparison between the non-covalently tethered reaction and the corresponding free/untethered reaction. It also avoids the complications that arise in a non-covalent three component kinase/substrate/scaffold system where there are additional bound and unbound states to consider (i.e. kinase and substrate segregated on different scaffold proteins).^24^ We expect that the observed effects from this model system should be generalizable to more complex multivalent assemblies.

Based on a minimal kinetic model, we predict that to observe a significant contribution from a tethered reaction, the substrate-scaffold interaction needs to be strong enough to appreciably bind at concentrations below the effective molarity (Scheme 1). In the minimal model, the substrate binds to the scaffold via a remote binding interaction and can then be phosphorylated in an intramolecular reaction. The free substrate can also bind directly to the active site of the kinase and react via an intermolecular reaction without engaging the remote binding site. The kinetic model indicates that the intramolecular, scaffold-mediated reaction (V_intra_) is faster than the intermolecular reaction (V_inter_) when the effective molarity is greater than *K*_D_ + [S], where *K*_D_ is the affinity of the substrate for the scaffold (Figure 4). If the affinity of the substrate for the scaffold is weak compared to the effective molarity (i.e. *K*_D_ > EM), there is no substrate concentration at which the intramolecular reaction is faster than the intermolecular reaction because the scaffolded complex does not significantly accumulate at concentrations below the effective molarity (Figure 4B). In contrast, if the affinity is strong relative to the effective molarity (i.e. *K*_D_ < EM, Figure 4C), then the intramolecular reaction will outcompete the intermolecular reaction at low substrate concentrations (i.e. when [S] < EM - *K*_D_). Even in this scenario, the intermolecular reaction will still outcompete the intramolecular reaction at sufficiently high substrate concentrations. The substrate concentration at which this switch occurs is determined by the effective molarity (Figure 4). Practically, our kinetic model suggests that we will observe the largest effects from tethering with a scaffold that recruits a substrate with a *K*_D_ < ∼0.08 µM, the effective molarity that we observed in the covalently-tethered kinase-substrate system (Figure 2). A similar conceptual analysis was recently described for tethered protein-ligand interactions.^25^

**Figure 4.**
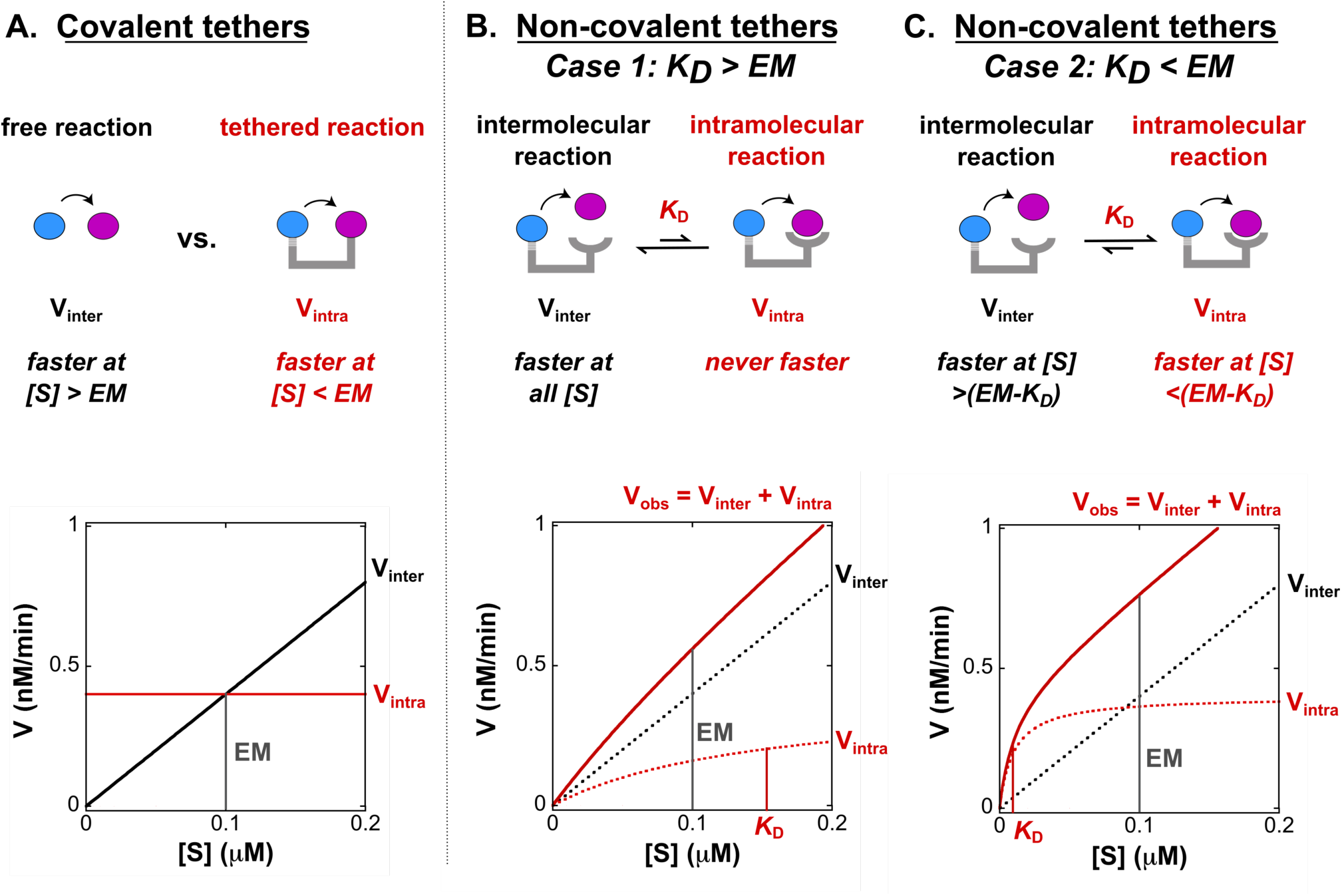
Predicted rates versus substrate concentration for various tethering strategies. (A) Observed rates vs. substrate concentration for a bimolecular reaction between kinase and free substrate (solid black line), and a covalently tethered kinase-substrate reaction (solid red line). The effective molarity (EM) is the substrate concentration at which V_inter_ and V_intra_ are equal (indicated by grey line). (B) Observed rates vs. substrate concentration for a model scaffold system where the remote tether binds to the substrate with a *K*_D_ > EM. There is no substrate concentration at which the rate of the intramolecular reaction (V_intra_, red dotted line) is faster than the intermolecular reaction (V_inter_, black dotted line). The observed rate (V_obs_, solid red line) will include contributions from both the intramolecular and intermolecular reactions (i.e. V_obs_ = V_intra_+V_inter_). (C) Observed rates vs. substrate concentration for a model scaffold system where the substrate binds to the remote tether with a *K*_D_ < EM. At low substrate concentrations (i.e. when [S] < EM - *K*_D_), the intramolecular reaction (V_intra_, red dotted line) is faster than the intermolecular reaction (V_inter_, black dotted line) because the complex assembles at concentrations below the EM. The total observed rate (V_obs_, solid red line) will include contributions from both the intramolecular and intermolecular reactions (i.e. V_obs_ = V_intra_+V_inter_). For all plots the effective molarity is indicated by a vertical grey line and is 0.1 µM. *K*_D_ values in B and C are indicated by a vertical red line and are 0.15 µM and 0.01 µM, respectively.

To construct a non-covalently tethered kinase substrate system, we used heterodimeric coiled-coil interactions to recruit the substrate to the kinase via a flexible tether (Figure 5A). We used the heterodimeric coiled-coils SYNZIP 5 and SYNZIP 6 because this pair is well-characterized structurally and biophysically and has been used for many applications in synthetic biology and protein design.^26-28^ We fused PKA to SYNZIP 6 (PKA-SYNZIP6) and the peptide substrate to SYNZIP5 (Pep-SYNZIP5). We measured a *K*_D_ of 0.01 µM for the Pep-SYNZIP5·PKA-SYNZIP6 interaction using fluorescence anisotropy (Figure S8); this binding affinity is consistent with previous measurements for SYNZIP5/6 affinity.^27^ Importantly, this affinity is below the predicted effective molarity of 0.08 µM, so we expect to observe a significant contribution from the tethered reaction at substrate concentrations <0.08 µM, but almost no effect on rates at substrate concentrations substantially above the effective molarity.

**Figure 5.**
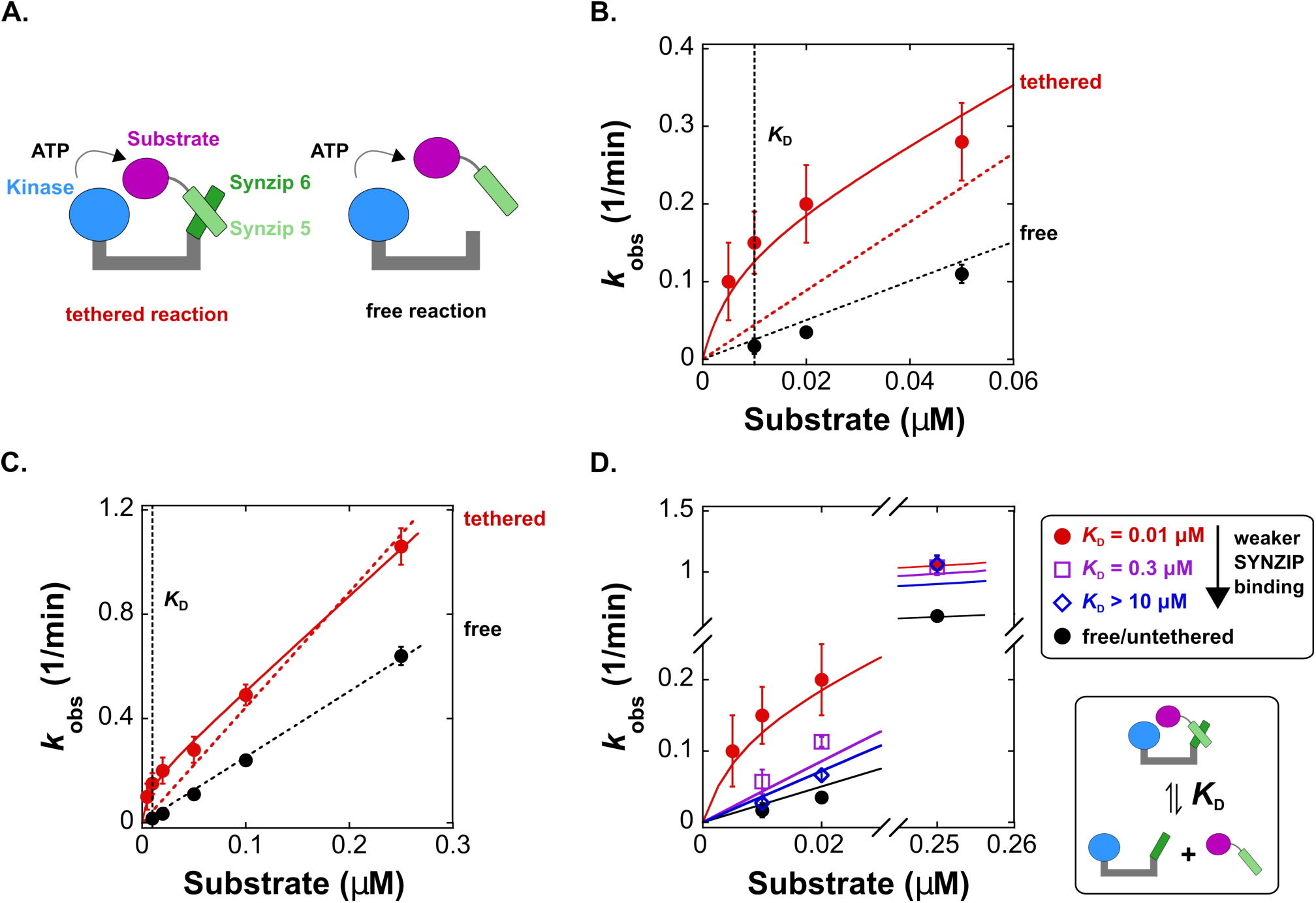
Non-covalent tethering enhances the rate of phosphorylation and depends on the relationship between effective molarity and *K*_D_. (A) Schematic of a model scaffold with a non-covalent tether that recruits a substrate to a kinase. The kinase is fused to SYNZIP6 (PKA-SYNZIP6) and the substrate is fused to SYNZIP5, which interact to form the tethered complex. In the corresponding free reaction, we used a kinase without SYNZIP6. (B) Plot of *k*_obs_ vs. [substrate] for the tethered and free reactions for [substrate] < 0.06 µM. At concentrations below the effective molarity, the values of *k*_obs_ for the tethered reaction are significantly larger than the untethered reaction. The solid red line is a fit to a kinetic model (Scheme 1 and Supporting Information) that includes a contribution from *k*_intra_, which accurately fits the tethered reaction data. The dotted lines are fits to a model where *k*_obs_ = (*k*_cat_/*K*_M_)[S] (this linear fit forces the line through zero). Both fits include data at higher substrate concentrations (full dataset is shown in panel C), and for the tethered reaction this linear model fails to account for the observed rate increase at low substrate concentrations. For the untethered reaction, the data scale linearly with [substrate]. The reaction of PKA-SYNZIP6 is also first order in enzyme concentration (Figure S9). (C) Plot of *k*_obs_ vs. [substrate] over an expanded range of substrate concentrations up to 0.25 µM. The dotted lines are fits to a simple linear model (*k*_obs_ = *k*_cat_/*K*_M_ [S]), and the solid red line is a fit to the complete model that includes *k*_intra_ (Scheme 1 and Supporting Information). For the untethered reaction, the slope corresponds to a bimolecular rate constant (*k*_cat_/*K*_M_) of 2.5 µM^-1^ min^-^ 1. For the tethered reaction, the value of *k*_cat_/*K*_M_ obtained from the fit to the complete model (solid red line) is 3.6 µM^-1^ min^-1^. This value differs from the untethered reaction by a factor of 1.4, likely due to prep-to-prep variations in the total activity of PKA-SYNZIP6 versus free PKA. (D) Plot of *k*_obs_ vs. [substrate] for the tethered reaction with the SYNZIP 5/6 pair and with two SYNZIP5 truncations that weaken the binding affinity (increasing *K*_D_) (Figure S10). At [substrate] <0.03 µM, values of *k*_obs_ decrease as *K*_D_ increases. At [substrate] = 0.25 µM, well above the effective molarity, values of *k*_obs_ are indistinguishable for substrates with different SYNZIP interaction affinities. The solid lines represent kinetic models using the values of *k*_intra_ and *k*_cat_/*K*_M_ from the untruncated pair in Figure 5C, and the corresponding *K*_D_ value from each truncated pair. Each [product] vs time trace was measured from separate reactions in duplicate. Error bars in B-D represent the standard error for *k*_obs_ obtained from a linear fit to [product] vs time for both datasets.

To test this prediction, we compared phosphorylation rates (*k*_obs_, where *k*_obs_ = V_obs_*/*[E]) for the tethered and free reactions over a range of substrate concentrations above and below the anticipated effective molarity. As expected, we observed a significant increase in *k*_obs_ for the tethered reaction, particularly at substrate concentrations <0.05 µM (Figure 5B). The observed values of *k*_obs_ for the tethered reaction are biphasic with [substrate] and correspond closely to the predictions of the kinetic model that includes contributions from an intramolecular reaction (Scheme 1). At low substrate concentrations, *k*_obs_ is dominated by contributions from the intramolecular reaction, while at high substrate concentrations *k*_obs_ is dominated by contributions from an intermolecular reaction of the tethered complex with free substrate (Figure 5B & C). For the untethered reaction, *k*_obs_ scaled linearly with substrate and enzyme concentrations, consistent with a bimolecular reaction (Figure 5B, C & Figure S9). Thus, tethering can enhance kinase reaction rates at substrate concentrations below the effective molarity.

Fitting the data to kinetic models allows us to calculate values for the effective molarity and define the conditions under which tethering produces significant rate enhancements. For the tethered reaction, the fit provides an intramolecular rate constant (*k*_intra_) of 0.15 min^-1^ for a substrate non-covalently tethered to enzyme (Figure 5B &C), similar to the *k*_intra_ value of 0.11 min^-1^ for covalently tethered substrate (Table 2). Using this *k*_intra_ value and the *k*_cat_/*K*_M_ value of 2.5 µM^-1^ min^-1^ derived from the free reaction (Figure 5B & C), we calculated an effective molarity of 0.06 µM for the non-covalently tethered system, similar to the effective molarity of 0.08 µM for the covalent system (Table 2). At substrate concentrations <0.05 µM (i.e. when [S] < EM - *K*_D_; EM = 0.06 µM and *K*_D_ = 0.01 µM), the substrate binds to the scaffold and primarily reacts with the kinase through the intramolecular, tethered reaction. Conversely, at substrate concentrations >0.05 µM (i.e. when [S] > EM - *K*_D_), the substrate predominantly reacts through a bimolecular pathway, even when PKA is fully occupied by a tethered substrate. Hence, we only observe significant effects from tethering with substrate concentrations less than ∼50 nM.

### A scaffold-substrate interaction that is weak relative to the EM does not accelerate the reaction

Our model predicts that observed rates from a tethered reaction should depend on the binding affinity between the substrate and the scaffold. If the affinity of the substrate for the scaffold is weak relative to the effective molarity (*K*_D_>>EM), then the intermolecular reaction is faster than the intramolecular reaction at all substrate concentrations (Figure 4B). As the binding affinity of the scaffold for the substrate increases, the model predicts that the intramolecular reaction will make increasingly smaller contributions to the observed rate.

To test this prediction, we made truncations of SYNZIP5 to weaken its affinity for SYNZIP6 and identified two substrates (Pep-SYNZIP5_trunc1_ and Pep-SYNZIP5_trunc2_) with binding affinities that were weaker than the cognate pair (0.3 µM and >10 µM, respectively) (Figure S10). We measured *k*_obs_ for PKA-SYNZIP6 with varying concentrations of truncated Pep-SYNZIP5 substrates. As expected, for substrate concentrations below the effective molarity (i.e. 0.01 or 0.02 µM), phosphorylation rates (*k*_obs_) decrease as *K*_D_ increases (Figure 5D). At a substrate concentration above the effective molarity (i.e. 0.25 µM), *k*_obs_ values are unaffected by binding affinity. These trends can be captured by our model using the values of *k*_intra_ and *k*_cat_/*K*_M_ derived from the cognate SYNZIP5/6 pair in Figure 5C, and the *K*_D_ value for each truncated pair. Moreover, this data confirms a key prediction from our model: as *K*_D_ increases, there will be less tethered complex assembled, which will therefore make a smaller contribution to observed rates. Thus, the relationship between the effective molarity and the *K*_D_ is a critical parameter that determines the magnitude of the contribution, if any, from a tethered reaction.

## Conclusions

Our results show that tethering a kinase and a substrate together can increase phosphorylation reaction rates. We show experimentally that the affinity for complex assembly plays a central role in the observed rates; if the affinity is weak compared to the effective molarity, a scaffolded complex will not assemble at concentrations where the intramolecular reaction is faster than the intermolecular reaction (Figure 5). This observation follows from a simple, intuitive kinetic model (Figure 4). Somewhat unexpectedly, however, the value of the effective molarity that we observe with a flexible scaffold model, ∼0.1 µM, is relatively small compared to effective molarities for small molecule reactions and protein ligand interactions.^12-14,22,23^ Natural protein-protein affinities span a broad range from low nM to high µM,^29,30^ and many scaffold-mediated interactions in signaling networks are likely to be well above a 0.1 µM effective molarity threshold.

If many biological interactions are too weak to bind below the effective molarity for a flexible scaffold, how do natural scaffold proteins make significant contributions to signaling networks? One possibility is that natural scaffold proteins achieve larger effective molarities with ordered structures that precisely orient kinase and substrate. Our understanding of scaffold protein structure remains limited, however, and many scaffold proteins are thought to contain intrinsically disordered regions.^29-31^ Future studies could test whether larger effective molarities can be obtained in engineered systems by using well-ordered, designed protein tethers. Alternatively, it is possible that flexible scaffolds have evolved relatively tight affinities, possibly via multivalent interactions, to ensure that complexes assemble in a concentration range where scaffold-mediated reactions can make a significant contribution to reaction rates. Finally, scaffold proteins can have many functions beyond tethering,^1,3,32,33^ and it may be that tethering-mediated rate enhancements are simply not the primary function of many scaffold proteins.

While many questions remain open, our work suggests two important conclusions with broad implications. First, many scaffold proteins have been defined by qualitative assays that indicate binding to a kinase and substrate. In the absence of quantitative data for reaction rates and binding constants, however, any inference that the function of these scaffold proteins is to accelerate reaction rates should be viewed with caution. Second, when attempts are made to engineer scaffolds in synthetic networks, careful consideration should be given to both linker structure and interaction affinities. Further studies on both natural and model scaffold proteins will help to further expand our understanding of complex signaling networks and improve our ability to engineer effective synthetic systems.

## Figures

**Scheme 1.**
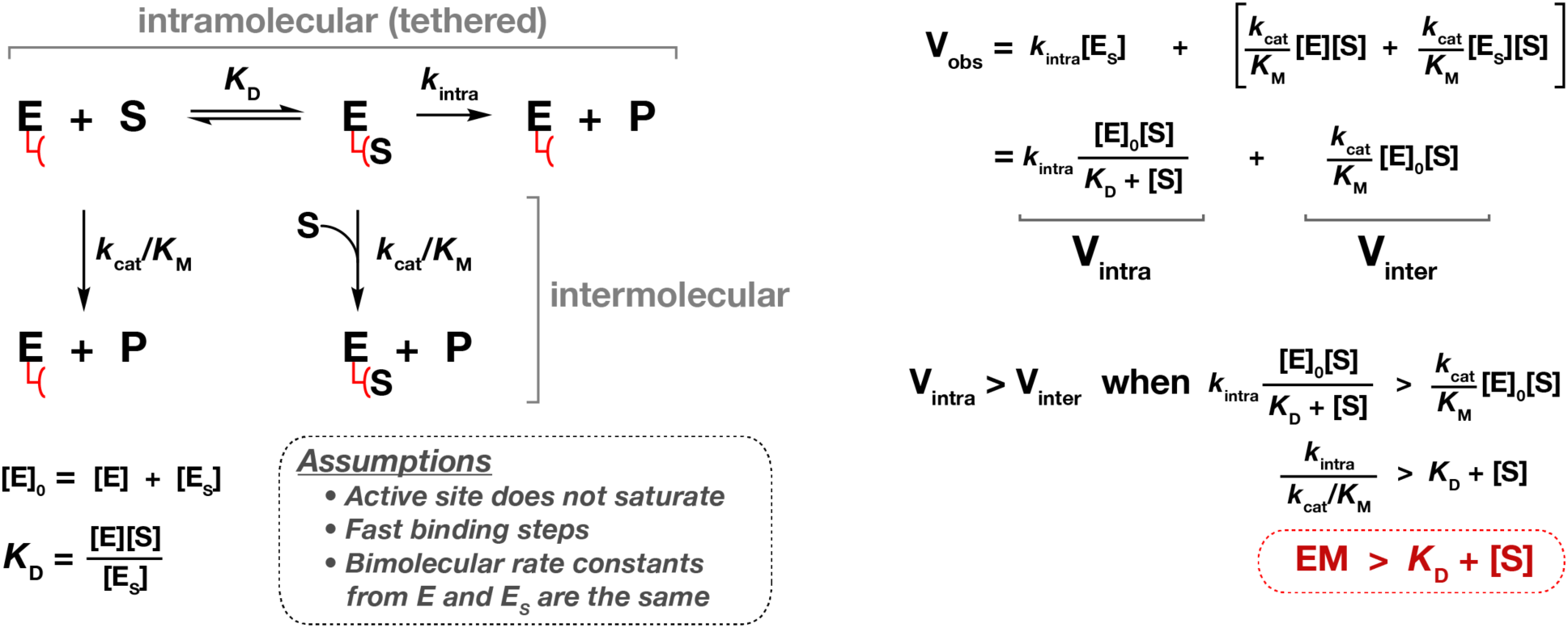
Minimal kinetic scheme for a noncovalent tethered reaction. The enzyme has a remote tether that can recruit a substrate, which then reacts with the active site through an intramolecular mechanism. Alternatively, the substrate can react with the enzyme through an intermolecular reaction by binding directly to the enzyme active site without engaging the tether. For simplicity, the intermolecular pathway does not include an E·S complex with substrate occupying the active site, because we did not detect saturation in any untethered reaction between PKA and the substrate peptide (Figure 2). We also assumed that the binding step to the tether is fast compared to *k*_intra_. Finally, we assumed that the bimolecular rate constants (*k*_cat_/*K*_M_) for E and E_S_ are the same. With these simplifying assumptions, the condition for when the intramolecular reaction (V_intra_) is faster than the intermolecular reaction (V_inter_) is: EM > *K*_D_ + [S]. See Supporting Information for a complete derivation.

## Acknowledgements

We thank Dustin Maly, Mike Gelb, Geeta Narlikar, Wendell Lim, Dan Herschlag, Maire Gavagan, and members of the Zalatan group for comments and discussion. This work was supported by a Career Award at the Scientific Interface from the Burroughs Wellcome Fund (J.G.Z.) and NIH R35 GM124773 (J.G.Z.). E.B.S. was funded by a Washington Research Foundation Innovation Postdoctoral Fellowship.

## Supporting Information

Supporting information, including protein sequences, supporting figures and a complete derivation of the equation shown in Scheme 1, are available free of charge.

## References

(1) Langeberg, L. K., and Scott, J. D. (2015) Signalling scaffolds and local organization of cellular behaviour. Nat. Rev. Mol. Cell Biol. 16, 232–244.

(2) Bhattacharyya, R. P., Reményi, A., Yeh, B. J., and Lim, W. A. (2006) Domains, motifs, and scaffolds: the role of modular interactions in the evolution and wiring of cell signaling circuits. Annu. Rev. Biochem. 75, 655–680.

(3) Good, M. C., Zalatan, J. G., and Lim, W. A. (2011) Scaffold proteins: hubs for controlling the flow of cellular information. Science 332, 680–686.

(4) Bashor, C. J., Helman, N. C., Yan, S., and Lim, W. A. (2008) Using engineered scaffold interactions to reshape MAP kinase pathway signaling dynamics. Science 319, 1539–1543.

(5) Dueber, J. E., Wu, G. C., Malmirchegini, G. R., Moon, T. S., Petzold, C. J., Ullal, A. V., Prather, K. L. J., and Keasling, J. D. (2009) Synthetic protein scaffolds provide modular control over metabolic flux. Nat. Biotechnol. 27, 753–759.

(6) Whitaker, W. R., Davis, S. A., Arkin, A. P., and Dueber, J. E. (2012) Engineering robust control of two-component system phosphotransfer using modular scaffolds. Proc. Natl. Acad. Sci. U.S.A. 109, 18090–18095.

(7) Park, S. H. (2003) Rewiring MAP Kinase Pathways Using Alternative Scaffold Assembly Mechanisms. Science 299, 1061–1064.

(8) Moon, J., and Park, S.-H. (2014) Reassembly of JIP1 Scaffold Complex in JNK MAP Kinase Pathway Using Heterologous Protein Interactions. PLoS One (Buday, L., Ed.) 9, e96797–7.

(9) Hobert, E. M., and Schepartz, A. (2012) Rewiring Kinase Specificity with a Synthetic Adaptor Protein. J. Am. Chem. Soc. 134, 3976–3978.

(10) Dueber, J. E., Yeh, B. J., Chak, K., and Lim, W. A. (2003) Reprogramming control of an allosteric signaling switch through modular recombination. Science 301, 1904–1908.

(11) Ryu, J., and Park, S.-H. (2015) Simple synthetic protein scaffolds can create adjustable artificial MAPK circuits in yeast and mammalian cells. Sci. Signal. 8, ra66–ra66.

(12) Kirby, A. J. (1980) Effective molarities for intramolecular reactions. Adv. Phys. Org. Chem. 17, 183–278.

(13) Krishnamurthy, V. M., Semetey, V., Bracher, P. J., Shen, N., and Whitesides, G. M. (2007) Dependence of Effective Molarity on Linker Length for an Intramolecular Protein-Ligand System. J. Am. Chem. Soc. 129, 1312–1320.

(14) Sørensen, C. S., and Kjaergaard, M. (2019) Effective concentrations enforced by intrinsically disordered linkers are governed by polymer physics. Proc. Natl. Acad. Sci. U.S.A. 116, 23124–23131.

(15) Das, A., Gerlits, O., Parks, J. M., Langan, P., Kovalevsky, A., and Heller, W. T. (2015) Protein Kinase A Catalytic Subunit Primed for Action: Time-Lapse Crystallography of Michaelis Complex Formation. Structure 23, 2331–2340.

(16) Zakeri, B., Fierer, J. O., Celik, E., Chittock, E. C., Schwarz-Linek, U., Moy, V. T., and Howarth, M. (2012) Peptide tag forming a rapid covalent bond to a protein, through engineering a bacterial adhesin. Proc. Natl. Acad. Sci. U.S.A. 109, E690–7.

(17) Picciotto, M. R., Cohn, J. A., Bertuzzi, G., Greengard, P., and Nairn, A. C. (1992) Phosphorylation of the cystic fibrosis transmembrane conductance regulator. J. Biol. Chem. 267, 12742–12752.

(18) Madhusudan, Trafny E. A., Xuong, N. H., Adams, J. A., Eyck, Ten L. F., Taylor, S. S., and Sowadski, J. M. (1994) cAMP-dependent protein kinase: crystallographic insights into substrate recognition and phosphotransfer. Protein Sci. 3, 176–187.

(19) Whitehouse, S., Feramisco, J. R., Casnellie, J. E., Krebs, E. G., and Walsh, D. A. (1983) Studies on the kinetic mechanism of the catalytic subunit of the cAMP-dependent protein kinase. J. Biol. Chem. 258, 3693–3701.

(20) Taylor, S. S., Knighton, D. R., Zheng, J., Sowadski, J. M., Gibbs, C. S., and Zoller, M. J. (1993) A template for the protein kinase family. Trends Biochem. Sci. 18, 84–89.

(21) Adams, J. A., and Taylor, S. S. (1993) Phosphorylation of peptide substrates for the catalytic subunit of cAMP-dependent protein kinase. J. Biol. Chem. 268, 7747–7752.

(22) Page, M. I., and Jencks, W. P. (1971) Entropic contributions to rate accelerations in enzymic and intramolecular reactions and the chelate effect. Proc. Natl. Acad. Sci. U.S.A. 68, 1678–1683.

(23) Van Valen, D., Haataja, M., and Phillips, R. (2009) Biochemistry on a Leash: The Roles of Tether Length and Geometry in Signal Integration Proteins. Biophys. J. 96, 1275–1292.

(24) Levchenko, A., Bruck, J., and Sternberg, P. W. (2000) Scaffold proteins may biphasically affect the levels of mitogen-activated protein kinase signaling and reduce its threshold properties. Proc. Natl. Acad. Sci. U.S.A. 97, 5818–5823.

(25) Sørensen, C. S., Jendroszek, A., and Kjaergaard, M. (2019) Linker Dependence of Avidity in Multivalent Interactions Between Disordered Proteins. J. Mol. Biol. 1–12.

(26) Reinke, A. W., Grant, R. A., and Keating, A. E. (2010) A Synthetic Coiled-Coil Interactome Provides Heterospecific Modules for Molecular Engineering. J. Am. Chem. Soc. 132, 6025–6031.

(27) Thompson, K. E., Bashor, C. J., Lim, W. A., and Keating, A. E. (2012) SYNZIP Protein Interaction Toolbox: *in Vitro* and *in Vivo* Specifications of Heterospecific Coiled-Coil Interaction Domains. ACS Synth. Biol. 1, 118–129.

(28) Park, W. M., Bedewy, M., Berggren, K. K., and Keating, A. E. (2017) Modular assembly of a protein nanotriangle using orthogonally interacting coiled coils. Sci. Rep. 7, 1–10.

(29) Smith, F. D., Reichow, S. L., Esseltine, J. L., Shi, D., Langeberg, L. K., Scott, J. D., and Gonen, T. (2013) Intrinsic disorder within an AKAP-protein kinase A complex guides local substrate phosphorylation. eLife 2, e01319.

(30) Noutsou, M., Duarte, A. M. S., Anvarian, Z., Didenko, T., Minde, D. P., Kuper, I., de Ridder, I., Oikonomou, C., Friedler, A., Boelens, R., Rüdiger, S. G. D., and Maurice, M. M. (2011) Critical Scaffolding Regions of the Tumor Suppressor Axin1 Are Natively Unfolded. J. Mol. Biol. 405, 773–786.

(31) Cortese, M. S., Uversky, V. N., and Keith Dunker, A. (2008) Intrinsic disorder in scaffold proteins: Getting more from less. Prog. Biophys. Mol. Biol. 98, 85–106.

(32) Nussinov, R., Ma, B., and Tsai, C.-J. (2013) A broad view of scaffolding suggests that scaffolding proteins can actively control regulation and signaling of multienzyme complexes through allostery. Biochim. Biophys. Acta 1834, 820–829.

(33) Mayer, B. J., Blinov, M. L., and Loew, L. M. (2009) Molecular machines or pleiomorphic ensembles: signaling complexes revisited. J. Biol. 8, 81.

(34) Good, M., Tang, G., Singleton, J., Reményi, A., and Lim, W. A. (2009) The Ste5 Scaffold Directs Mating Signaling by Catalytically Unlocking the Fus3 MAP Kinase for Activation. Cell 136, 1085–1097.

