## Supporting Information for "The relationship between effective molarity and affinity governs rate enhancements in tethered kinase-substrate reactions"

Elizabeth B. Speltz and Jesse G. Zalatan
Department of Chemistry, University of Washington, Seattle, WA 98195, USA

*Corresponding author: Jesse Zalatan


**This PDF file includes:**

Figures S1 to S11
Protein Sequences
Complete derivation of the kinetic model in Scheme 1
Supplemental References

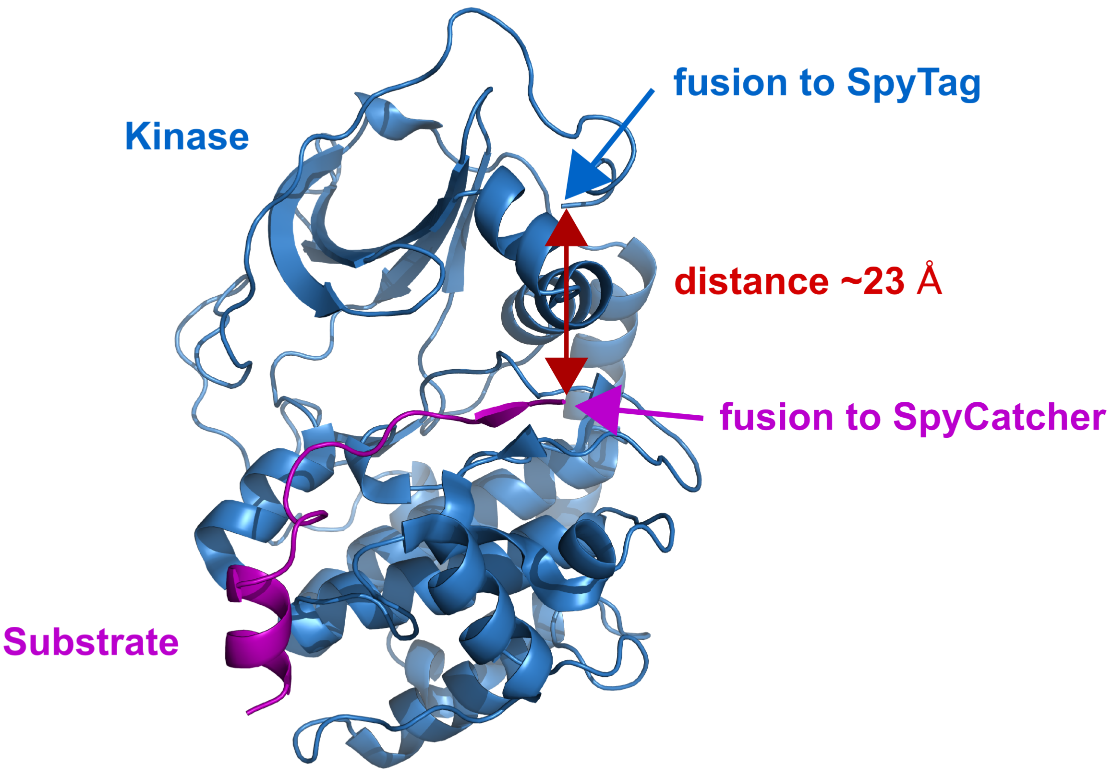


**Figure S1. Crystal structure of a PKA-substrate complex.**
Crystal structure of the catalytic subunit of PKA (blue) bound to a peptide substrate with sequence TTYADFIASGRTGRRASIHD (purple).[1] In our model system for a covalently-tethered kinase-substrate complex, the C-terminus of PKA is fused to SpyTag and the C-terminus of a peptide substrate is fused to SpyCatcher as indicated by arrows. The distance between these C-termini is ~23 Å. Note that the peptide substrate used in our model system (sequence FGEKRKNSILNPI) extends the C-terminus of the peptide shown in the crystal structure by an additional two residues. Thus, the crystal structure provides only a rough estimate for the distance between SpyTag and SpyCatcher fusions. The structure was rendered in PyMol using PDB ID 3X2U.

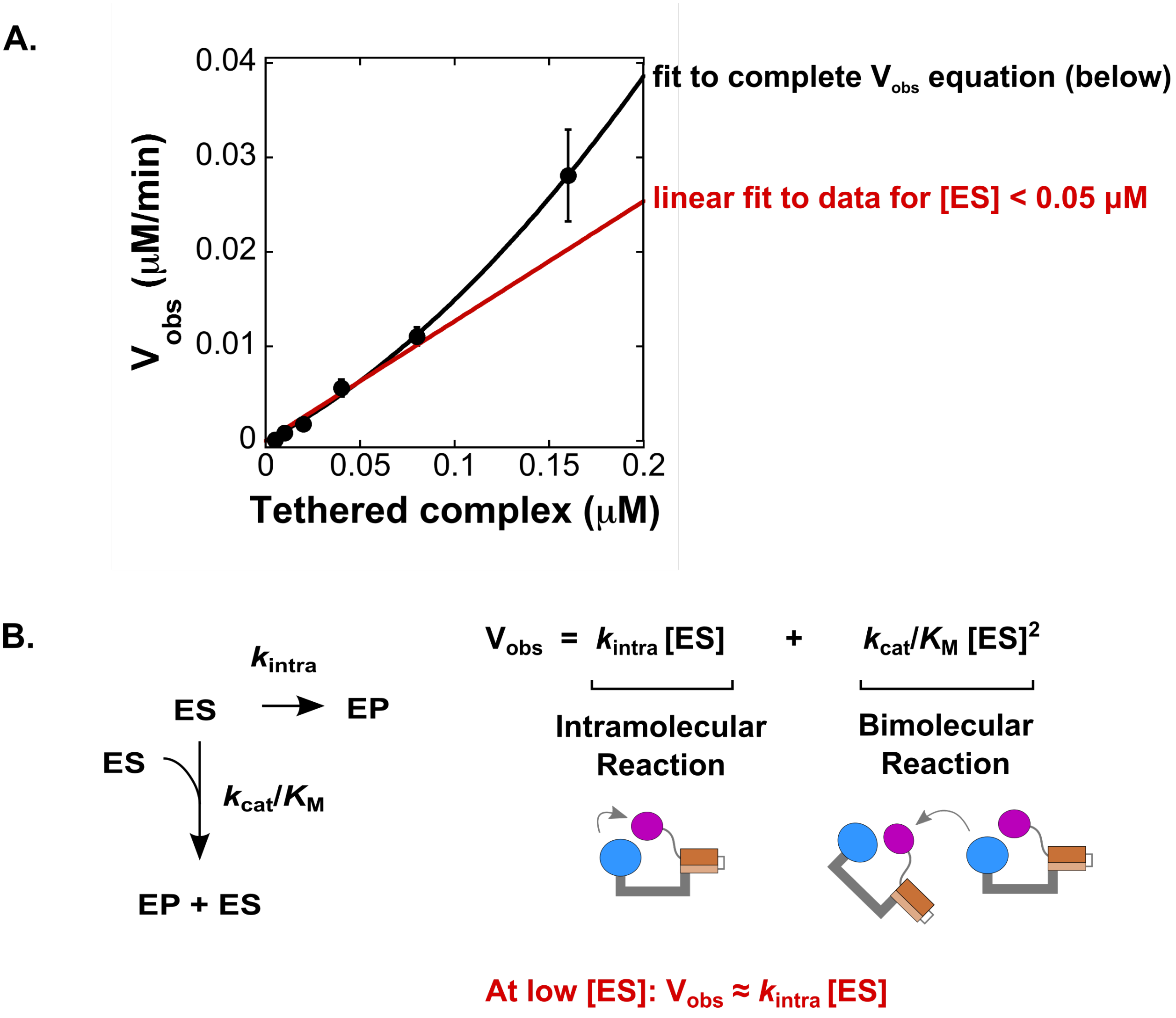


**Figure S2. Measurement of $k_{intra}$ for covalently tethered complexes.**
A. Plot of $V_{obs}$ vs [tethered complex] for the covalently tethered complex with a four-residue linker between the substrate and SpyCatcher. To extract $k_{intra}$, the data is fit to the complete $V_{obs}$ equation shown in part B (black line). Alternatively, the data at low [ES] (<50 nM) can be fit to a line, which corresponds to $V_{obs} = k_{intra}$[ES] (red line).
B. Definition of $V_{obs}$ for a covalently tethered enzyme-substrate (ES) complex. The ES complex can react via an intramolecular reaction ($k_{intra}$), or two ES complexes can react in a bimolecular reaction ($k_{cat}/K_M$). EP is the tethered enzyme-product complex. The observed rate includes contributions from both the intramolecular and bimolecular reactions. At low concentrations of ES, the equation can be approximated as $V_{obs}= k_{intra}$ [ES].

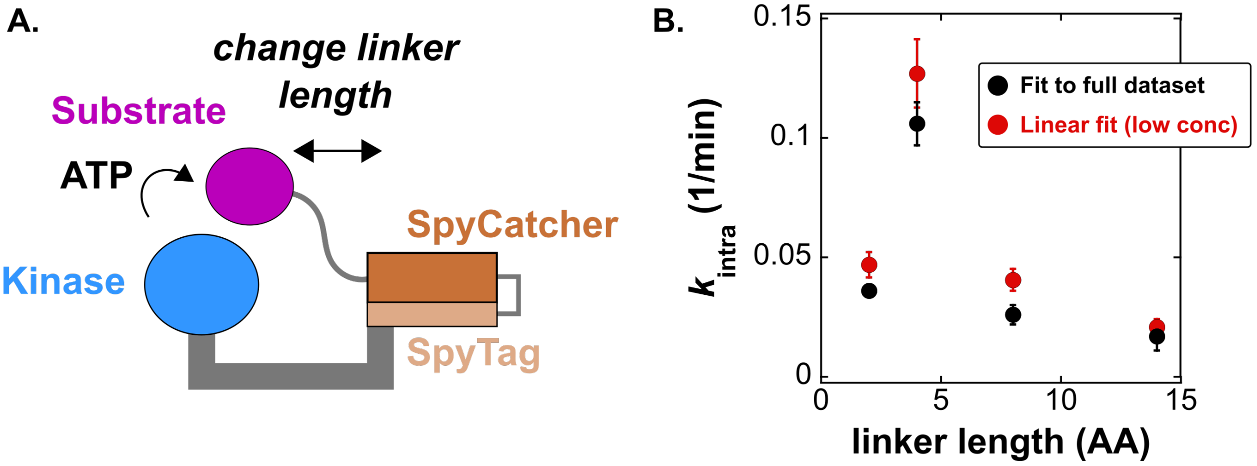


**Figure S3. Comparison of $k_{intra}$ for covalently tethered complexes.**

A. Schematic of the covalently-tethered complex. The SpyCatcher-SpyTag complex covalently links the kinase with its substrate. We varied the length of the linker that connects the substrate to SpyCatcher from two to fourteen residues and measured the unimolecular rate constant ($k_{intra}$).

B. $k_{intra}$ was measured by fitting the linear portion of the data at low [tethered complex] (red circles) or by fitting the full $V_{obs}$ dataset (black circles – as shown in main text Fig 2). For details and derivations for both fitting methods, refer to Fig S2. Both fitting methods produce values of $k_{intra}$ that are in close agreement. Each [product] vs time trace was measured from separate reactions in duplicate. Error bars represent the standard error for $k_{intra}$ obtained from the fits to [product] vs time for both datasets.

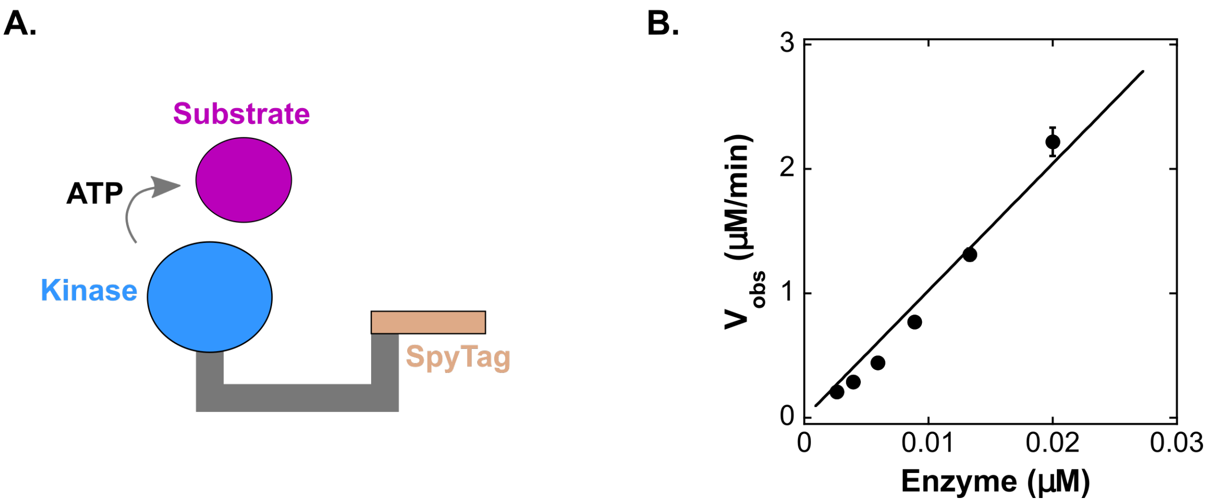


**Figure S4. $V_{obs}$ scales linearly with enzyme concentration for PKA-SpyTag.**
(A) Schematic of the reaction between PKA-SpyTag and substrate. We measured initial rates ($V_{obs}$) for varying concentrations of PKA-SpyTag and a saturating concentration of a model substrate (150 uM kemptide).[2]
(B) Plot of $V_{obs}$ vs. [PKA-SpyTag]. $V_{obs}$ is linear over all PKA-SpyTag concentrations tested, indicating that the observed reaction is first order in enzyme. Together with the observed first order kinetics with substrate (Figure 3A), these data indicate that the reaction is bimolecular. Each [product] vs time trace was measured from separate reactions in duplicate. Error bars represent the standard error for $k_{obs}$ obtained from a linear fit to [product] vs time for both datasets.

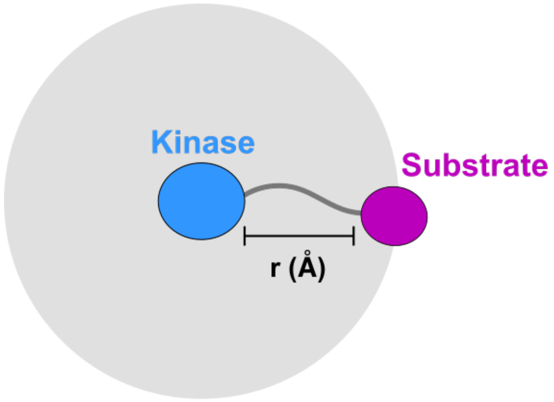


$$r\ (\text{Å}) = \text{linker length (aa)} * 3.8\ (\text{Å/aa})$$

$$\text{Effective Concentration (M)} = \frac{1}{4/3\ \pi\ r^{3}}\ \frac{10^{27}\ \text{Å}^{3}}{\text{L}}\ \frac{1\ \text{mol}}{6.022 \times 10^{23}\ \text{molecules}}$$

**Figure S5. Estimation of effective concentration for a covalently tethered substrate.** The distance between the kinase and the substrate (r) is represented by the number of amino acids in the linker. To calculate the radius for the covalently-tethered kinase-substrate complex, we assumed that the total linker length between the kinase and the substrate is represented by the sum of the linker length between the kinase and SpyTag (8 aa) and the linker length between the substrate and SpyCatcher (4 aa). Thus, the total linker length is 12 aa. Using a length per aa value of 3.8 Å/aa, this simplified model predicts an effective concentration of ~4 mM.[3]

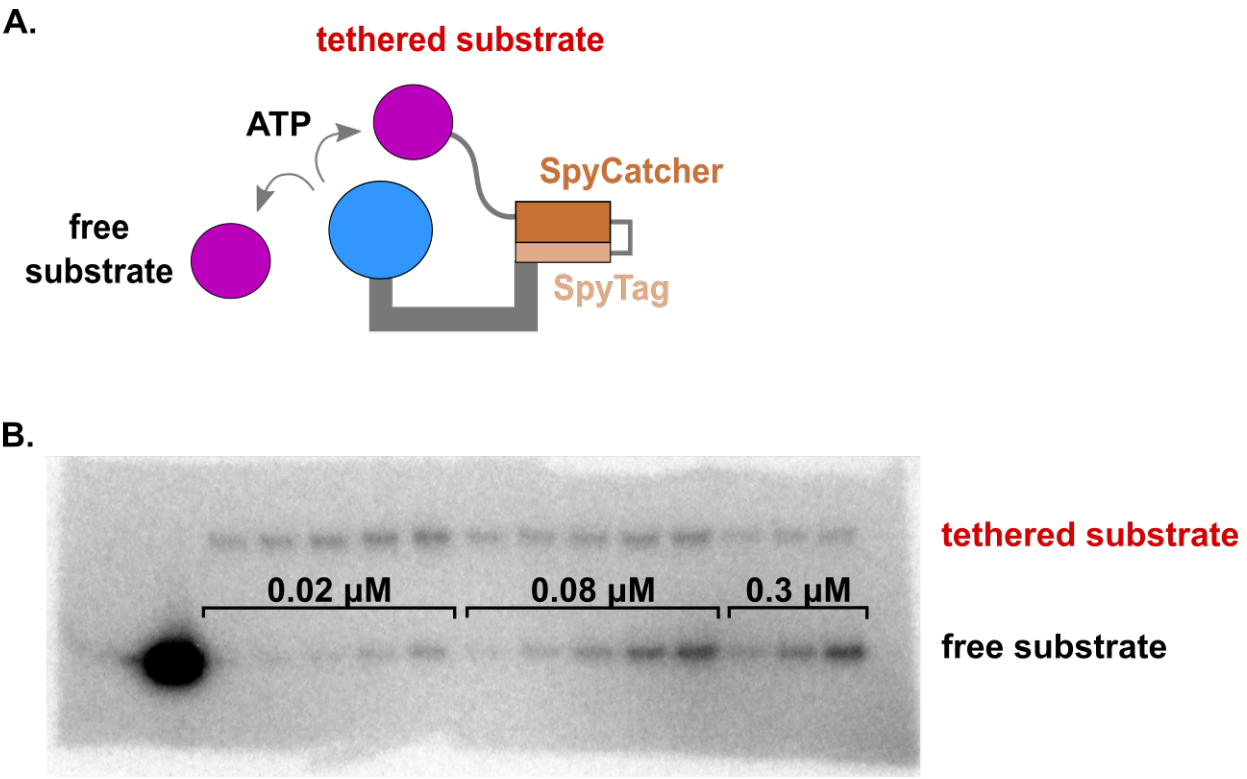


**Fig S6. Competition experiment between tethered and free substrate.**
A. Schematic representation of the components in the competition assay. Tethered PKA-substrate is mixed with varying concentrations of free substrate. Phosphorylation of the tethered and free substrate are monitored simultaneously by SDS-PAGE and autoradiography.
B. Representative gel from a competition experiment. The top band represents the phosphorylated tethered product and the bottom band represents the phosphorylated free product. The concentration of the free substrate is indicated above each lane.

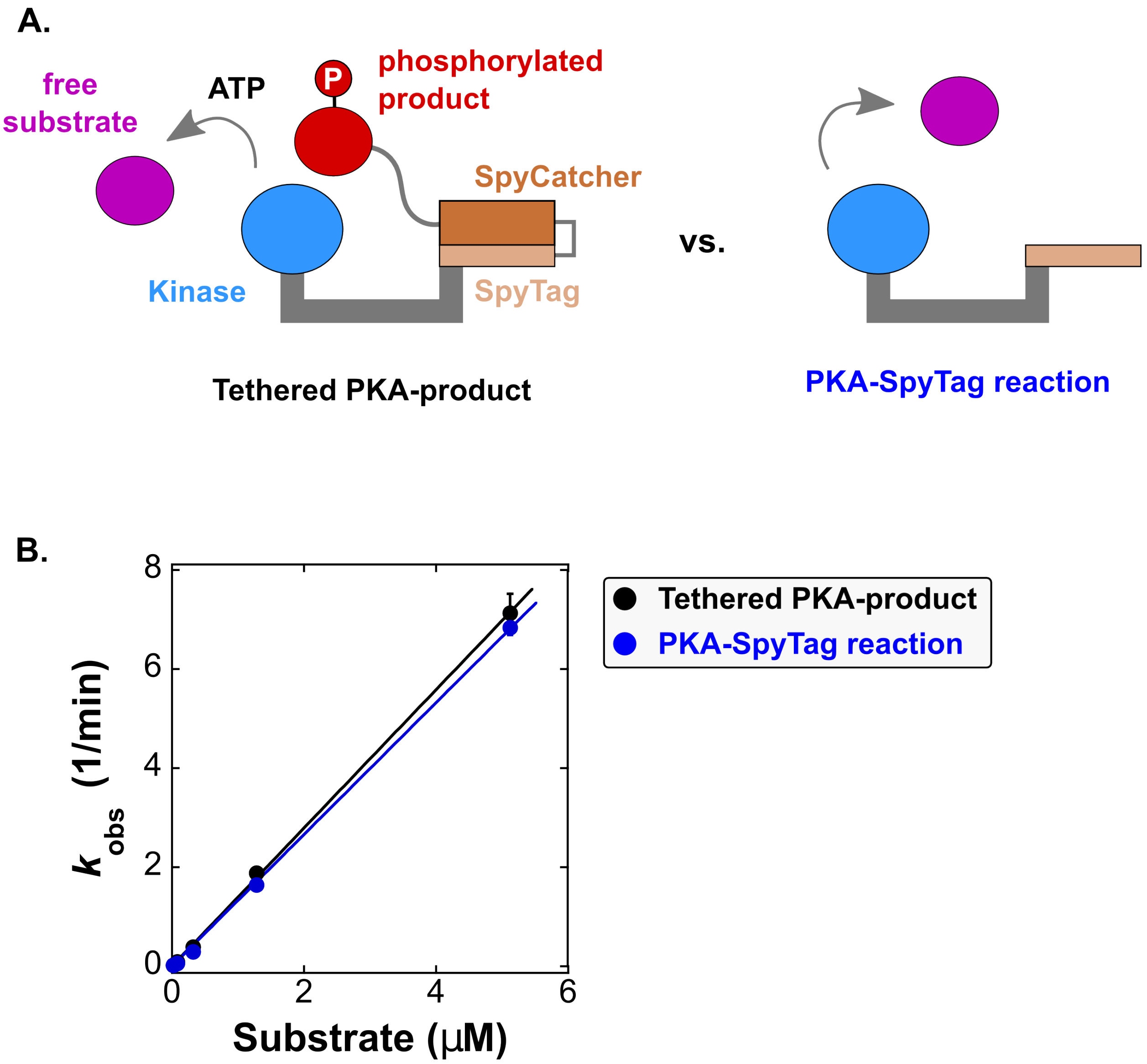


**Figure S7. Covalently tethered enzyme is not inhibited by product.**
A. Schematic of the reaction between a tethered PKA-product complex and free substrate compared to the reaction between untethered PKA (PKA-SpyTag) and free substrate. To make the tethered PKA-product complex, we allowed the four-residue linker tethered PKA-substrate complex to react to completion. Briefly, we incubated 2.5 μM tethered PKA-substrate complex with 250 μM ATP for two hours at room temperature. To measure reaction rates, the fully phosphorylated complex was diluted to 2.5 nM and mixed with 100 μM cold ATP containing 0.02 μCi/ μL γ-$^{32}$P-ATP. Both the untethered PKA-SpyTag and PKA-phosphosubstrate complex reactions were initiated by the addition of free substrate.
B. Plot of $k_{obs}$ vs [substrate] for PKA-SpyTag with free substrate (blue) or a PKA-phosphosubstrate complex with free substrate (black). The slope of this data corresponds to the bimolecular rate constant ($k_{cat}/K_M$) and is indistinguishable in these reactions: 1.3 min$^{-1}$ μM$^{-1}$ for PKA-SpyTag with free substrate and 1.4 min$^{-1}$ μM$^{-1}$ for the PKA-phosphosubstrate complex with free substrate.

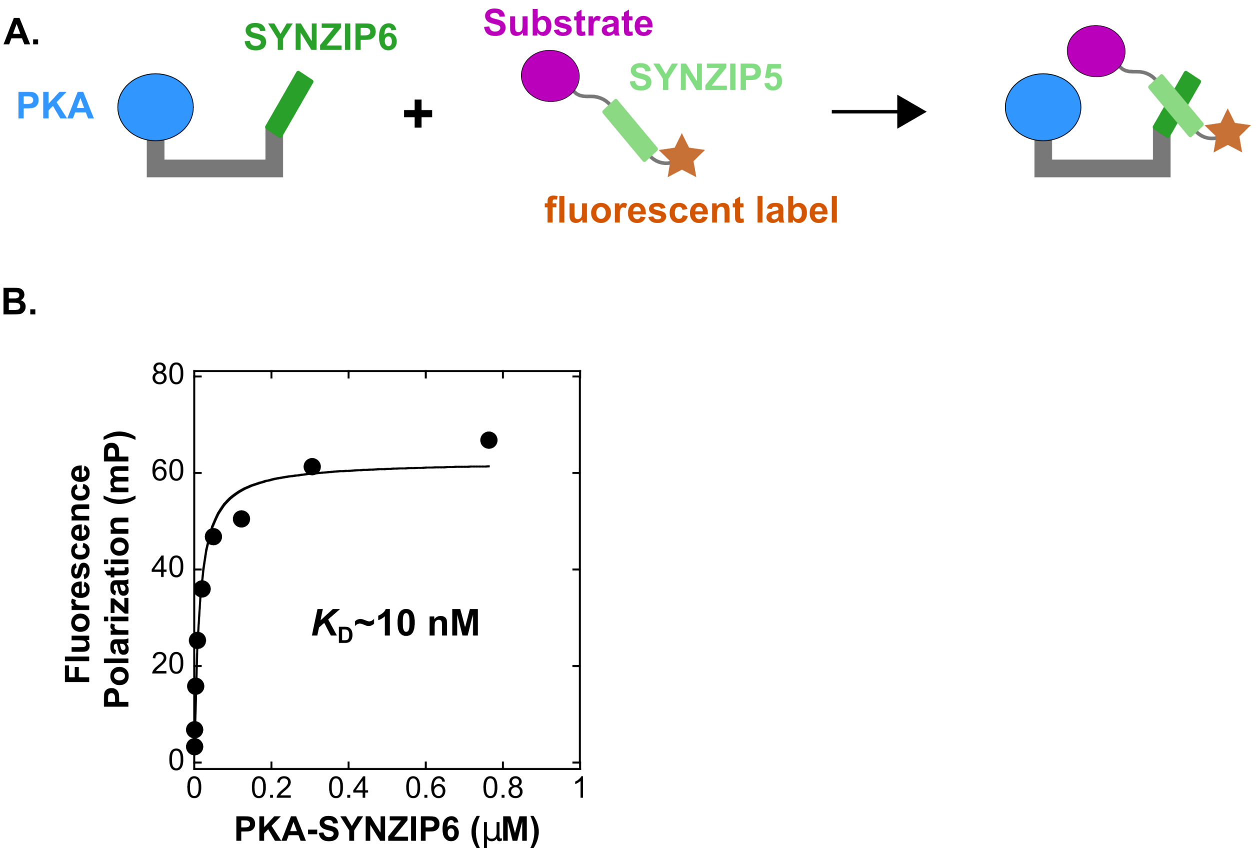


**Fig S8. Measurement of the binding affinity between Pep-SYNZIP5 and PKA-SYNZIP6.**
A. Schematic of the fluorescence polarization assay. We monitored the fluorescence polarization of Pep-SYNZIP5* (Pep-SYNZIP5 labeled with BODIPY-TAMRA) and PKA-SYNZIP6 as described previously.[4] As PKA-SYNZIP6 binds to Pep-SYNZIP5*, fluorescence polarization of the dye on the substrate increases.
B. Plot of fluorescence polarization as a function of [PKA-SYNZIP6]. The data is fit to a 1:1 binding model, which gives a $K_D$ of 10 nM. Data points are the average of two replicates.

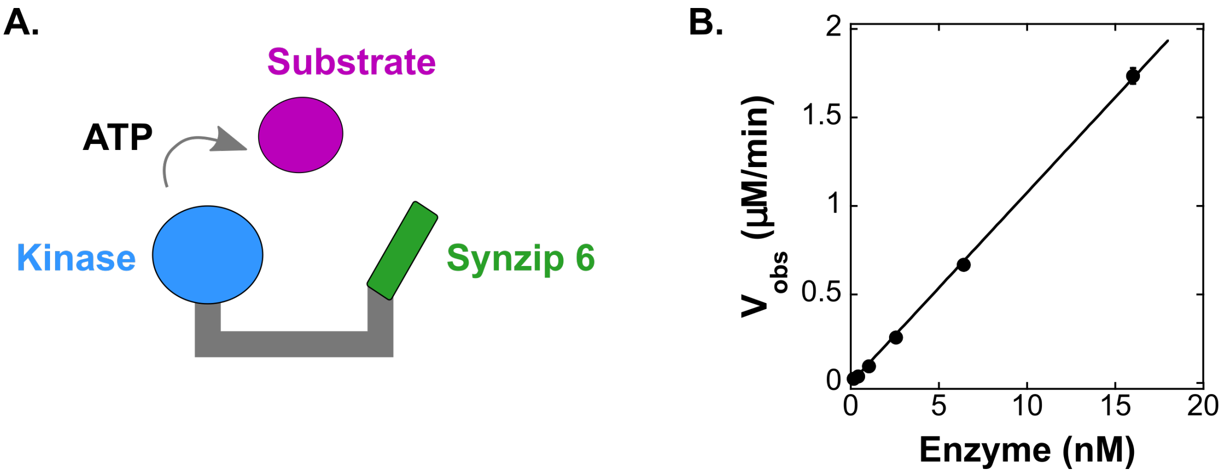


**Figure S9. $V_{obs}$ scales linearly with enzyme concentration for PKA-SYNZIP6.**
(A) Schematic of the reaction between PKA-SYNZIP6 and substrate. We measured initial rates ($V_{obs}$) for varying concentrations of PKA-SYNZIP6 and a saturating concentration of a model substrate (150 uM kemptide).[2]
(B) Plot of $V_{obs}$ vs. [PKA-SYNZIP6]. $V_{obs}$ is linear over all PKA-SYNZIP6 concentrations tested, indicating that the observed reaction is first order in enzyme. Together with the observed first order kinetics with substrate (Figure 5C), these data indicate that the reaction is bimolecular. Each [product] vs time trace was measured from separate reactions in duplicate. Error bars represent the standard error for $k_{obs}$ obtained from a linear fit to [product] vs time for both datasets.

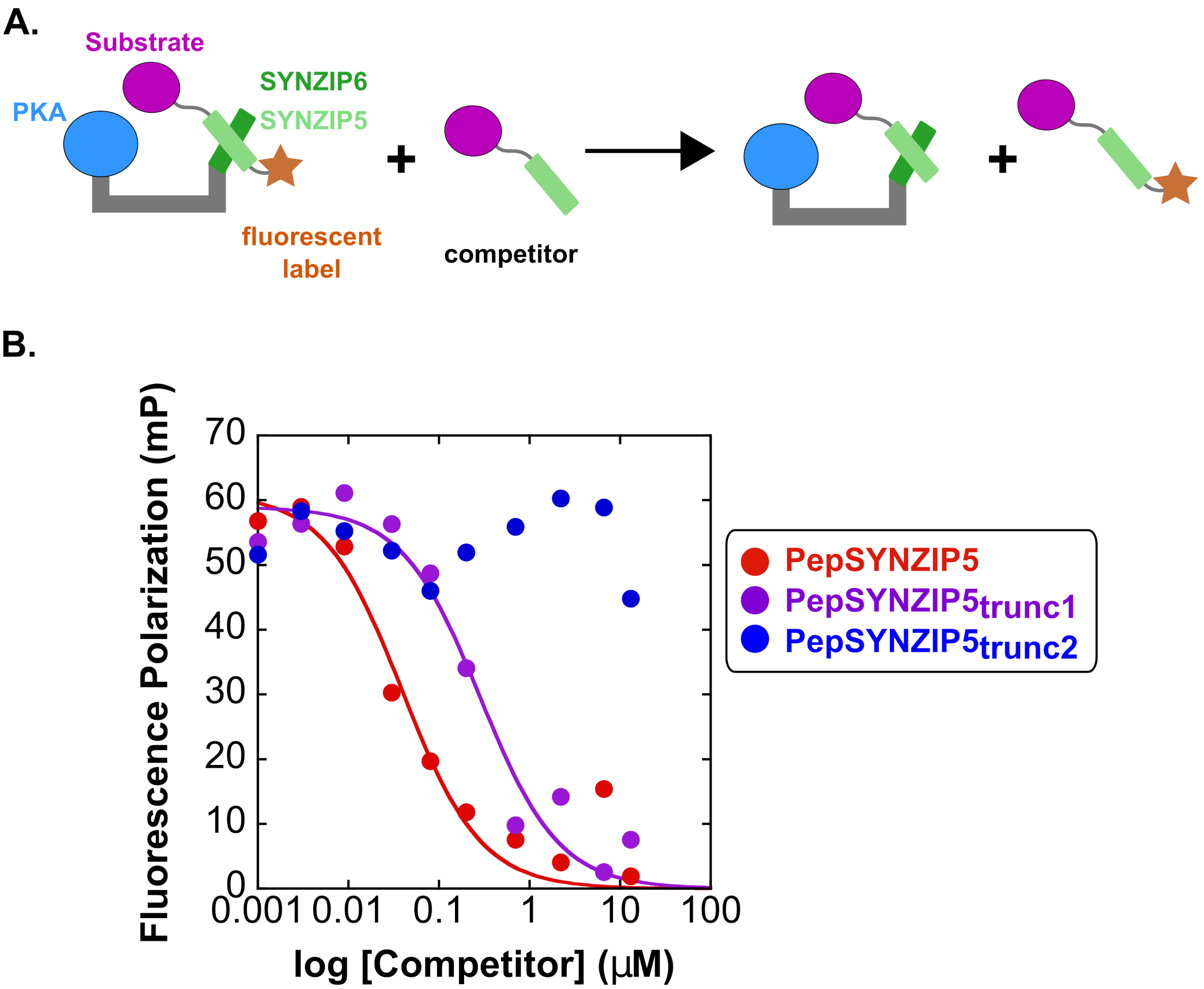


**Fig S10. Binding constants for truncated SYNZIP5 constructs from competition anisotropy measurements.**

A. Schematic of the competition binding assay. 1 nM Pep-SYNZIP5* (Pep-SYNZIP5 labeled with BODIPY-TAMRA) and 30 nM PKA-SYNZIP6 are mixed with varying concentrations of an unlabeled substrate competitor. As the competitor binds to PKA-SYNZIP6, the fluorescence polarization of the dye on Pep-SYNZIP5* will decrease.

B. Plot of fluorescence polarization vs. [competitor] (semi-log scale). The background polarization from non-specific binding of competitor and labeled substrate is subtracted from all measurements. The data is fit to a 1:1 binding model (see methods). The $K_D$ values are as follows: PepSYNZIP5 = 0.01 μM, PepSYNZIP5$_{trunc1}$ = 0.3 μM, PepSYNZIP5$_{trunc2}$ >10 μM. Data points are the average of two replicates.

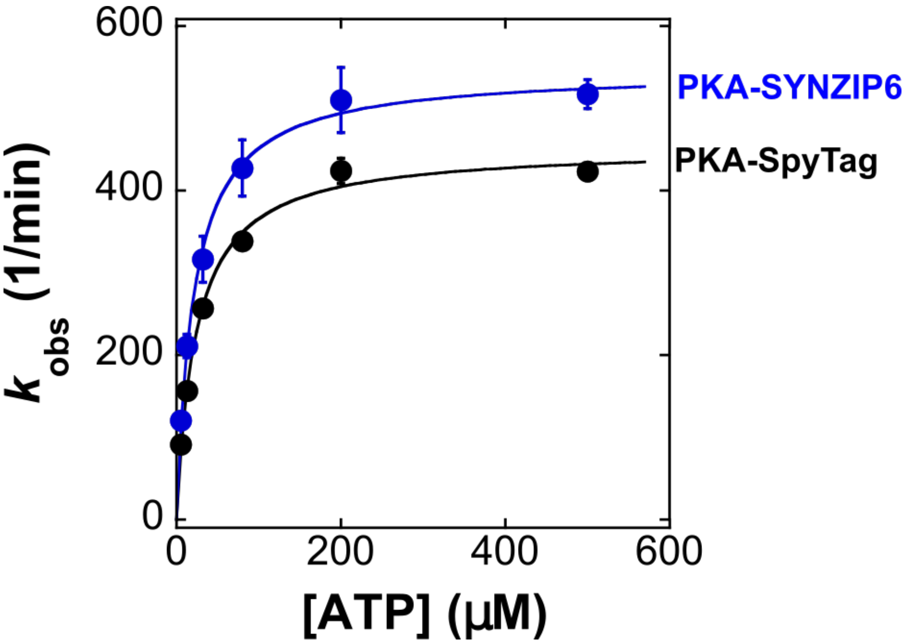


**Figure S11. Measurement of $K_M$ for ATP for PKA constructs.**
Plot of $k_{obs}$ vs. [ATP] for PKA-SYNZIP6 (blue dots) and PKA-SpyTag (black dots) with 150 μM substrate and varying [ATP]. For these experiments, we used a well-characterized peptide substrate (Kemptide) because it can saturate the PKA active site.[2] To obtain the $K_M$ for ATP, the data is fit to the Michaelis-Menten equation. For PKA-SYNZIP6, the $K_M$ is 24 μM; for PKA-SpyTag, the $K_M$ is 21 μM. Each [product] vs time trace was measured from separate reactions in duplicate. Error bars represent the standard error for $k_{obs}$ obtained from a linear fit to [product] vs time for both datasets.

**Protein Sequences**

Annotations:
PKA
SpyTag
SpyCatcher
Peptide substrate
SYNZIP 6
SYNZIP 5

**pES263 pMAL PKA 8x SpyTag**

MKIEEGKLVIWINGDKGYNGLAEVGKKFEKDTGIKVTVEHPDKLEEKFPQVAATGDGPDIIFWA
HDRFGGYAQSGLLAEITPDKAFQDKLYPFTWDAVRYNGKLIAYPIAVEALSLIYNKDLLPNPPK
TWEEIPALDKELKAKGKSALMFNLQEPYFTWPLIAADGGYAFKYENGKYDIKDVGVDNAGAKAG
LTFLVDLIKNKHMNADTDYSIAEAAFNKGETAMTINGPWAWSNIDTSKVNYGVTVLPTFKGQPS
KPFVGVLSAGINAASPNKELAKEFLENYLLTDEGLEAVNKDKPLGAVALKSYEEELAKDPRIAA
TMENAQKGEIMPNIPQMSAFWYAVRTAVINAASGRQTVDEALKDAQTNSSSNNNNNNNNNNLGI
EGRISTSGSGGGGGSMSENLYFQGSSVKEFLAKAKEDFLKKWETPSQNTAQLDQFDRIKTLGTG
SFGRVMLVKHKESGNHYAMKILDKQKVVKLKQIEHTLNEKRILQAVNFPFLVKLEFSFKDNSNL
YMVMEYVAGGEMFSHLRRIGRFSEPHARFYAAQIVLTFEYLHSLDLIYRDLKPENLLIDQQGYI
QVTDFGFAKRVKGRTWTLCGTPEYLAPEIILSKGYNKAVDWWALGVLIYEMAAGYPPFFADQPI
QIYEKIVSGKVRFPSHFSSDLKDLLRNLLQVDLTKRFGNLKNGVNDIKNHKWFATTDWIAIYQR
KVEAPFIPKFKGPGDTSNFDDYEEEEIRVSINEKCGKEFTEFTSGGSGGTAHIVMVDAYKPTKS
GRHHHHHH

**pES367 CFTR 2x SpyCatcher**

MGHHHHHHDYDIPTTENLYFQGSFGEKRKNSILNPIGTDSATHIKFSKRDEDGKELAGATMELR
DSSGKTISTWISDGQVKDFYLYPGKYTFVETAAPDGYEVATAITFTVNEQGQVTVNGKATKGDA
HI

**pES368 CFTR 4x SpyCatcher**

MGHHHHHHDYDIPTTENLYFQGSFGEKRKNSILNPISGGTDSATHIKFSKRDEDGKELAGATME
LRDSSGKTISTWISDGQVKDFYLYPGKYTFVETAAPDGYEVATAITFTVNEQGQVTVNGKATKG
DAHI

**pES264 CFTR 8x SpyCatcher**

MGHHHHHHDYDIPTTENLYFQGSFGEKRKNSILNPITSGGSGGTDSATHIKFSKRDEDGKELAG
ATMELRDSSGKTISTWISDGQVKDFYLYPGKYTFVETAAPDGYEVATAITFTVNEQGQVTVNGK
ATKGDAHI

**pES343 CFTR 14x SpyCatcher**

MGHHHHHHDYDIPTTENLYFQGSFGEKRKNSILNPIGGSGGSGGGTGGPGDSATHIKFSKRDED
GKELAGATMELRDSSGKTISTWISDGQVKDFYLYPGKYTFVETAAPDGYEVATAITFTVNEQGQ
VTVNGKATKGDAHI

**pES044 pMAL CFTR**

MKIEEGKLVIWINGDKGYNGLAEVGKKFEKDTGIKVTVEHPDKLEEKFPQVAATGDGPDIIFWA
HDRFGGYAQSGLLAEITPDKAFQDKLYPFTWDAVRYNGKLIAYPIAVEALSLIYNKDLLPNPPK
TWEEIPALDKELKAKGKSALMFNLQEPYFTWPLIAADGGYAFKYENGKYDIKDVGVDNAGAKAG
LTFLVDLIKNKHMNADTDYSIAEAAFNKGETAMTINGPWAWSNIDTSKVNYGVTVLPTFKGQPS
KPFVGVLSAGINAASPNKELAKEFLENYLLTDEGLEAVNKDKPLGAVALKSYEEELAKDPRIAA
TMENAQKGEIMPNIPQMSAFWYAVRTAVINAASGRQTVDEALKDAQTNSSSNNNNNNNNNNLGI
EGRISTSGSGGGGGSMSENLYFQGSFGEKRKNSILNPIPGSGRHHHHHH

**pES234 pMAL PKA 8x SYNZIP6**

MKIEEGKLVIWINGDKGYNGLAEVGKKFEKDTGIKVTVEHPDKLEEKFPQVAATGDGPDIIFWA
HDRFGGYAQSGLLAEITPDKAFQDKLYPFTWDAVRYNGKLIAYPIAVEALSLIYNKDLLPNPPK
TWEEIPALDKELKAKGKSALMFNLQEPYFTWPLIAADGGYAFKYENGKYDIKDVGVDNAGAKAG
LTFLVDLIKNKHMNADTDYSIAEAAFNKGETAMTINGPWAWSNIDTSKVNYGVTVLPTFKGQPS
KPFVGVLSAGINAASPNKELAKEFLENYLLTDEGLEAVNKDKPLGAVALKSYEEELAKDPRIAA
TMENAQKGEIMPNIPQMSAFWYAVRTAVINAASGRQTVDEALKDAQTNSSSNNNNNNNNNNLGI
EGRISTSGSGGGGGSMSENLYFQGSSVKEFLAKAKEDFLKKWETPSQNTAQLDQFDRIKTLGTG
SFGRVMLVKHKESGNHYAMKILDKQKVVKLKQIEHTLNEKRILQAVNFPFLVKLEFSFKDNSNL
YMVMEYVAGGEMFSHLRRIGRFSEPHARFYAAQIVLTFEYLHSLDLIYRDLKPENLLIDQQGYI
QVTDFGFAKRVKGRTWTLCGTPEYLAPEIILSKGYNKAVDWWALGVLIYEMAAGYPPFFADQPI
QIYEKIVSGKVRFPSHFSSDLKDLLRNLLQVDLTKRFGNLKNGVNDIKNHKWFATTDWIAIYQR
KVEAPFIPKFKGPGDTSNFDDYEEEEIRVSINEKCGKEFTEFTSGGSGGTQKVAQLKNRVAYKL
KENAKLENIVARLENDNANLEKDIANLEKDIANLERDVARSGRHHHHHH

**pES311 pMAL CFTR 14x SYNZIP 5**

MKIEEGKLVIWINGDKGYNGLAEVGKKFEKDTGIKVTVEHPDKLEEKFPQVAATGDGPDIIFWA
HDRFGGYAQSGLLAEITPDKAFQDKLYPFTWDAVRYNGKLIAYPIAVEALSLIYNKDLLPNPPK
TWEEIPALDKELKAKGKSALMFNLQEPYFTWPLIAADGGYAFKYENGKYDIKDVGVDNAGAKAG
LTFLVDLIKNKHMNADTDYSIAEAAFNKGETAMTINGPWAWSNIDTSKVNYGVTVLPTFKGQPS
KPFVGVLSAGINAASPNKELAKEFLENYLLTDEGLEAVNKDKPLGAVALKSYEEELAKDPRIAA
TMENAQKGEIMPNIPQMSAFWYAVRTAVINAASGRQTVDEALKDAQTNSSSNNNNNNNNNNLGI
EGRISTSGSGGGGGSMSENLYFQGSFGEKRKNSILNPIGGSGGSGGGTGGPGNTVKELKNYIQE
LEERNAELKNLKEHLKFAKAELEFELAAHKFESGRHHHHHH

**pES346 pMAL CFTR 14x SYNZIP5 trunc1**

MKIEEGKLVIWINGDKGYNGLAEVGKKFEKDTGIKVTVEHPDKLEEKFPQVAATGDGPDIIFWA
HDRFGGYAQSGLLAEITPDKAFQDKLYPFTWDAVRYNGKLIAYPIAVEALSLIYNKDLLPNPPK
TWEEIPALDKELKAKGKSALMFNLQEPYFTWPLIAADGGYAFKYENGKYDIKDVGVDNAGAKAG
LTFLVDLIKNKHMNADTDYSIAEAAFNKGETAMTINGPWAWSNIDTSKVNYGVTVLPTFKGQPS

KPFVGVLSAGINAASPNKELAKEFLENYLLTDEGLEAVNKDKPLGAVALKSYEEELAKDPRIAA
TMENAQKGEIMPNIPQMSAFWYAVRTAVINAASGRQTVDEALKDAQTNSSSNNNNNNNNNNLGI
EGRISTSGSGGGGGSMSENLYFQGSFGEKRKNSILNPIGGSGGSGGGTGGPGNTVKELKNYIQE
LEERNAELKNLKEHLKFAKAELESGRHHHHHH

**pES349 pMAL CFTR 14x SYNZIP5 trunc2**

MKIEEGKLVIWINGDKGYNGLAEVGKKFEKDTGIKVTVEHPDKLEEKFPQVAATGDGPDIIFWA
HDRFGGYAQSGLLAEITPDKAFQDKLYPFTWDAVRYNGKLIAYPIAVEALSLIYNKDLLPNPPK
TWEEIPALDKELKAKGKSALMFNLQEPYFTWPLIAADGGYAFKYENGKYDIKDVGVDNAGAKAG
LTFLVDLIKNKHMNADTDYSIAEAAFNKGETAMTINGPWAWSNIDTSKVNYGVTVLPTFKGQPS
KPFVGVLSAGINAASPNKELAKEFLENYLLTDEGLEAVNKDKPLGAVALKSYEEELAKDPRIAA
TMENAQKGEIMPNIPQMSAFWYAVRTAVINAASGRQTVDEALKDAQTNSSSNNNNNNNNNNLGI
EGRISTSGSGGGGGSMSENLYFQGSFGEKRKNSILNPIGGSGGSGGGTGGPGNTVKELKNYIQE
LEERNAELKNLKESGRHHHHHH

### Derivation of the kinetic model from Scheme 1

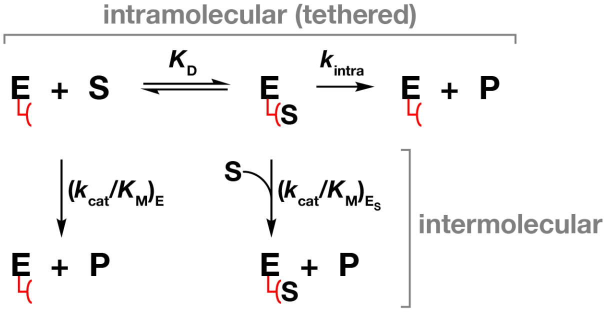


$$V_{intra} > V_{inter} \text{ when } k_{intra}\frac{[E]_0[S]}{K_D + [S]} > \frac{k_{cat}}{K_M}[E]_0[S]$$

$$\frac{k_{intra}}{k_{cat}/K_M} > K_D + [S]$$

$$EM > K_D + [S]$$

*Assumptions*
- *Active site does not saturate*
- *Fast binding steps*

**This model suggests that the intramolecular reaction is faster than the intermolecular reaction only when the scaffolded complex assembles at concentrations below the effective molarity (i.e. EM > $K_D$).**

1) Define relationship between total enzyme ($E_0$), free enzyme (E), and tethered enzyme-substrate ($E_S$):

$$[E]_0 = [E] + [E_S]$$

2) Define relationship between E and $E_S$:
**Using the equilibrium constant implicitly assumes that binding to tether is fast compared to $k_{intra}$*

$$K_D = \frac{[E][S]}{[E_S]}$$

3) Relate $E_0$ to $E_S$:

$$[E]_0 = [E_S]\frac{K_D}{[S]} + [E_S] = [E_S]\left(\frac{K_D}{[S]} + 1\right)$$

4) Relate $E_0$ to E:

$$[E]_0 = [E] + [E]\frac{[S]}{K_D} = [E]\left(1 + \frac{[S]}{K_D}\right)$$

5) Define $V_{obs}$ in terms of each possible pathway to product:
**$V_{inter}$ terms assume no significant buildup of an E·S complex with substrate occupying the active site. The observed rate is represented by a bimolecular term.*

$$\begin{aligned} V_{obs} &= V_{intra} + V_{inter} \\ &= V_{intra} + \left[V_{inter\,(E)} + V_{inter\,(E_S)}\right] \\ &= k_{intra}[E_S] + \left[\left(\frac{k_{cat}}{K_M}\right)_E [E][S] + \left(\frac{k_{cat}}{K_M}\right)_{E_S} [E_S][S]\right] \end{aligned}$$

6) Use (3) to substitute $[E]_0$ in place of $[E_S]$ in the $V_{intra}$ term:

$$V_{obs} = k_{intra} \frac{[E]_0}{\left(\frac{K_D}{[S]} + 1\right)} + \left[\left(\frac{k_{cat}}{K_M}\right)_E [E][S] + \left(\frac{k_{cat}}{K_M}\right)_{E_S} [E_S][S]\right]$$

$$= k_{intra} \frac{[E]_0[S]}{(K_D + [S])} + \left[\left(\frac{k_{cat}}{K_M}\right)_E [E][S] + \left(\frac{k_{cat}}{K_M}\right)_{E_S} [E_S][S]\right]$$

7) Assume the bimolecular rate constants from E and $E_S$ are the same:
(i.e. $\left(\frac{k_{cat}}{K_M}\right)_E$ and $\left(\frac{k_{cat}}{K_M}\right)_{E_S}$ are equal) - s*ee (9) below for the case where this assumption is not made.*

$$V_{obs} = k_{intra} \frac{[E]_0[S]}{(K_D + [S])} + \left[\left(\frac{k_{cat}}{K_M}\right)_E [E][S] + \left(\frac{k_{cat}}{K_M}\right)_{E_S} [E_S][S]\right]$$

$$= k_{intra} \frac{[E]_0[S]}{(K_D + [S])} + \frac{k_{cat}}{K_M}([E] + [E_S])[S]$$

$$= k_{intra} \frac{[E]_0[S]}{(K_D + [S])} + \frac{k_{cat}}{K_M}[E]_0[S]$$

$$k_{obs} = k_{intra} \frac{[S]}{(K_D + [S])} + \frac{k_{cat}}{K_M}[S]$$

8) Identify the conditions that allow the intramolecular reaction to be faster than the intermolecular reaction. Specifically, in the case where $V_{intra} > V_{inter}$, determine the relationship between effective molarity (EM), $K_D$, and [S]:

If $V_{intra} > V_{inter}$, then:

$$k_{intra} \frac{[E]_0[S]}{(K_D + [S])} > \frac{k_{cat}}{K_M}[E]_0[S]$$

$$\frac{k_{intra}}{k_{cat}/K_M} > K_D + [S]$$

$$EM > K_D + [S]$$

Interpretation: the effective molarity (EM) must be greater than ($K_D$ + [S]) for the intramolecular reaction to be faster than the intermolecular reaction.

9) If $\left(\frac{k_{cat}}{K_M}\right)_E$ and $\left(\frac{k_{cat}}{K_M}\right)_{E_S}$ are <u>not</u> equal, then use (3) and (4) to substitute $[E]_0$ for [E] and $[E_S]$:

$$
\begin{aligned}
V_{obs} &= k_{intra}\frac{[E]_0[S]}{(K_D+[S])} + \left[\left(\frac{k_{cat}}{K_M}\right)_E [E][S] + \left(\frac{k_{cat}}{K_M}\right)_{E_S}[E_S][S]\right] \\
&= k_{intra}\frac{[E]_0[S]}{(K_D+[S])} + \left[\left(\frac{k_{cat}}{K_M}\right)_E \frac{[E]_0}{\left(1+{[S]}/{K_D}\right)} + \left(\frac{k_{cat}}{K_M}\right)_{E_S}\frac{[E]_0}{\left({K_D}/{[S]}+1\right)}\right][S] \\
&= k_{intra}\frac{[E]_0[S]}{(K_D+[S])} + \left[\left(\frac{k_{cat}}{K_M}\right)_E \frac{1}{\left(1+{[S]}/{K_D}\right)} + \left(\frac{k_{cat}}{K_M}\right)_{E_S}\frac{1}{\left({K_D}/{[S]}+1\right)}\right][E]_0[S] \\
&= k_{intra}\frac{[E]_0[S]}{(K_D+[S])} + k_{intra\,(obs)}[E]_0[S]
\end{aligned}
$$

$$
k_{intra\,(obs)} = \left[\left(\frac{k_{cat}}{K_M}\right)_E \frac{1}{\left(1+{[S]}/{K_D}\right)} + \left(\frac{k_{cat}}{K_M}\right)_{E_S}\frac{1}{\left({K_D}/{[S]}+1\right)}\right]
$$

When [S] >> $K_D$: $k_{intra\,(obs)} = \left(\frac{k_{cat}}{K_M}\right)_{E_S}$

When [S] << $K_D$: $k_{intra\,(obs)} = \left(\frac{k_{cat}}{K_M}\right)_E$

### Supporting References

(1) Das, A., Gerlits, O., Parks, J. M., Langan, P., Kovalevsky, A., and Heller, W. T. (2015) Protein Kinase A Catalytic Subunit Primed for Action: Time-Lapse Crystallography of Michaelis Complex Formation. *Structure 23*, 2331–2340.
(2) Kemp, B. E., Graves, D. J., Benjamini, E., and Krebs, E. G. (1977) Role of multiple basic residues in determining the substrate specificity of cyclic AMP-dependent protein kinase. *J. Biol. Chem. 252*, 4888–4894.
(3) Bertagna, A., Toptygin, D., Brand, L., and Barrick, D. (2008) The effects of conformational heterogeneity on the binding of the Notch intracellular domain to effector proteins: a case of biologically tuned disorder. *Biochem. Soc. Trans. 36*, 157–166.
(4) Thompson, K. E., Bashor, C. J., Lim, W. A., and Keating, A. E. (2012) SYNZIP Protein Interaction Toolbox: *in Vitro* and *in Vivo* Specifications of Heterospecific Coiled-Coil Interaction Domains. *ACS Synth. Biol. 1*, 118–129.
